# The ISPpu9 insertion sequence of *Pseudomonas putida* KT2440 generates various circular intermediates enabling modular transposition

**DOI:** 10.1101/2025.01.17.633520

**Authors:** Elena Parés-Guillén, Luis Yuste, Fernando Rojo, Renata Moreno

## Abstract

*Pseudomonas putida* KT2440 contains seven copies of an insertion sequence (IS) belonging to the IS*110* family, designated ISPpu9, which are inserted into repetitive extragenic palindromic sequences. Five of these copies include a small RNA, *ssr9*, located downstream of the transposase gene. Additionally, three separate copies of *ssr9* are also present. When transferred to a different *P. putida* strain, ISPpu9 inserted at a specific target site highly similar to that of the KT2440 strain. Ssr9 was not needed for transposition. Circular intermediates were detected containing either the transposase, the transposase along with *ssr9*, or *ssr9* alone. This finding explains the presence of these distinct modules in KT2440 chromosome. Minicircle formation required the transposase and short sequences located within the IS ends, which were identified. Unlike other ISs of this family, minicircle formation in ISPpu9 did not produce a hybrid promoter to enhance transcription of the ISPpu9 transposase. Instead, transcription occurred efficiently from its native promoter. In contrast, minicircles formed by ISPpu10, another IS of the same family also present in KT2440, generated a hybrid promoter significantly stronger than its weak native transposase promoter. This suggests that hybrid promoters might only emerge when the native transposase promoter is inherently weak.

## INTRODUCTION

Insertion Sequences (ISs) are small mobile genetic elements that include only the components required for transposition and a regulatory element to control transposition frequency (1). They are grouped into families according to their sequence similarity to other ISs and their transposition mechanism (2). Most ISs exhibit low or no sequence specificity for target selection, although some show a preference, or even specificity, for precise sites with particular DNA sequences or secondary structures. Upon insertion, ISs often generate a short direct repeat at both ends of the target sequence.

ISs play an important role in the evolution of bacterial genomes, as they can generate gene mutations, deletions or inversions of large DNA regions (1,3,4). In addition, ISs can generate composite transposons that facilitate the movement of genes, including virulence or antibiotic-resistance genes. ISs are therefore important for bacterial adaptation to their environment.

*Pseudomonas putida* KT2440 is a soil bacterium that has become a model for biotechnological applications due to its lack of virulence, resistance to harsh conditions, and versatile metabolism (5,6,7). The genome of this strain, 6.2 Mb in length (8), contains 36 IS elements, most of them present in multiple copies (9). One of these ISs, named ISPpu9, is present as seven copies scattered throughout the chromosome and inserted at a specific target located within short conserved sequences belonging to the so-called repetitive <u>e</u>xtragenic <u>p</u>alindromic (REP) family (8,10; see Fig. 1A). REPs are highly conserved sequences containing an imperfect palindrome, found mostly in non-coding (extragenic) regions. Many bacterial species contain REPs, although with a different sequence in each case (11,12,13). *P. putida* KT2440 has more than 800 REPs with a conserved 35 bp sequence (12). REP sequences serve as targets for some IS elements (10,14).

**Figure 1.**
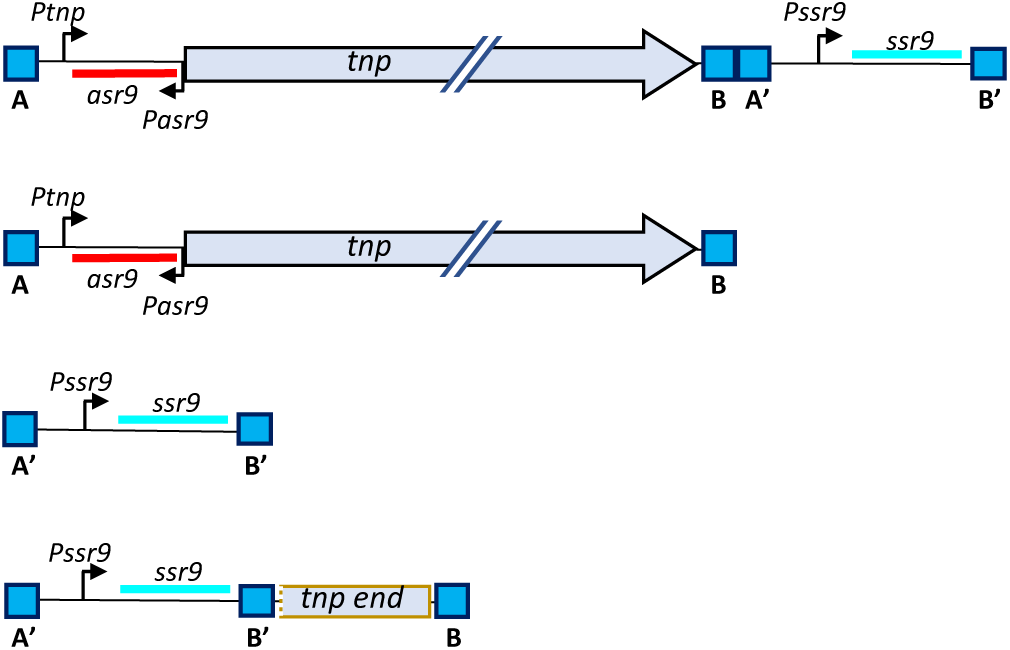
ISPpu9 variants found in *P. putida* KT1440 genome. This strain contains seven copies of the ISPpu9 transposase gene (*tnp*), all associated with the *asr9* gene encoding the Asr9 sRNA (in red) and flanked by conserved A and B boxes. Downstream of 5 of these *tnp* copies, the *ssr9* gene encoding the Ssr9 sRNA (in blue) is present, flanked by A’ and B’ boxes, which differ from the A and B boxes by 1-3 nt (see Table S2, Supplementary Material). The remaining two *tnp* copies lack an associated *ssr9* gene. Additionally, three extra copies of *ssr9* are located at distinct genome sites, all flanked by A’ and B’ boxes. One of these (between the *rlmB* and *rpsF* genes) includes a fragment of the 3’ end of *tnp* followed by a B box. Promoters for *asr9*, *tnp* and *ssr9* are indicated with arrows.

ISPpu9 contains a single ORF that codes for a DEDD-type transposase of the IS*110*/IS*492* family (Fig. 1B). These transposases differ from those of other families in that the N-terminal domain shows homology with RuvC Holliday junction resolvases, suggesting a distinctive mechanism of action (reviewed in ref. 1). The IS*110*/IS*492* family can be divided in two groups, the IS*110* group, which includes ISs with no inverted repeats at their left and right ends, and the IS*1111* group, which show sub-terminal inverted repeat sequences at their ends (1,15,16). ISPpu9 belongs to the IS*110* group which includes ISs known to insert at specific DNA sites, such as IS*492* from *Pseudoalteromonas atlantica* (17,18) and IS*621* from *Escherichia coli* (19). In several members of the IS*110* and IS*1111* families, transposition has been found to generate a double-stranded circular intermediate (20,21,15,16). This has also been observed in some ISs of the IS*3*, IS*21* and IS*30* families (reviewed in 22). Joining of the right and left ends of the IS can generate a new promoter, named *Pjunc*, which includes a -35 region provided by one IS end and a -10 region from the other end, at the proper distance. This promoter can drive transcription of the transposase gene, increasing its expression (20–23). The *Pjunc* promoter generated at the minicircles of the IS*110* and IS*1111* family members also drives the expression of a small non-coding RNA that binds to the transposase and facilitates recognition of the target site (15, 16). This RNA, named bridge RNA (15) or seek RNA (16), is typically found upstream of the transposase gene in the IS*110* group members, and transcribed in the same direction, whereas it is located downstream of the transposase in the IS*1111* group members. The bridge RNA binds to the transposase and includes two loops containing short nucleotide patches that can base-pair with complementary sequences at the donor and target DNAs, respectively, thus facilitating an RNA-guided DNA recombination event (15,16,24).

*P. putida* KT2440 contains two variants of ISPpu9 (Fig. 1). The first includes the transposase (*tnp*) and the antisense Asr9 sRNA, which is believed to stimulate *tnp* mRNA translation (25). The second variant has an additional sRNA downstream of the *tnp* gene, named Ssr9 and which might counteract the activating effect of Asr9 (25). The variant lacking *ssr9* is flanked by sequences that are also found flanking the *ssr9* sRNA gene; those flanking the first variant have been named “A” and “B”, while those flanking *ssr9* have herein been named A’ and B’ to differentiate them from the former ones (Fig. 1; sequence presented in Fig. 2A). The A and B boxes are 43 bp and 16 bp long, respectively, including the terminal AG dinucleotide present at the 5’ end of the A box and at the 3’ end of the B box which defines the boundaries of the IS. The genome of strain KT2440 contains all three possible combinations: three copies of an AB’ segment containing the transposase, *asr9* and *ssr9*, two copies of an AB segment containing *tnp* and *asr9* only, and three copies of an A’B’ segment containing *ssr9* only. This suggests that excision and insertion of ISPpu9 occurs by binding of the transposase to the A/A’ and B/B’ sites in all three possible combinations.

**Figure 2.**
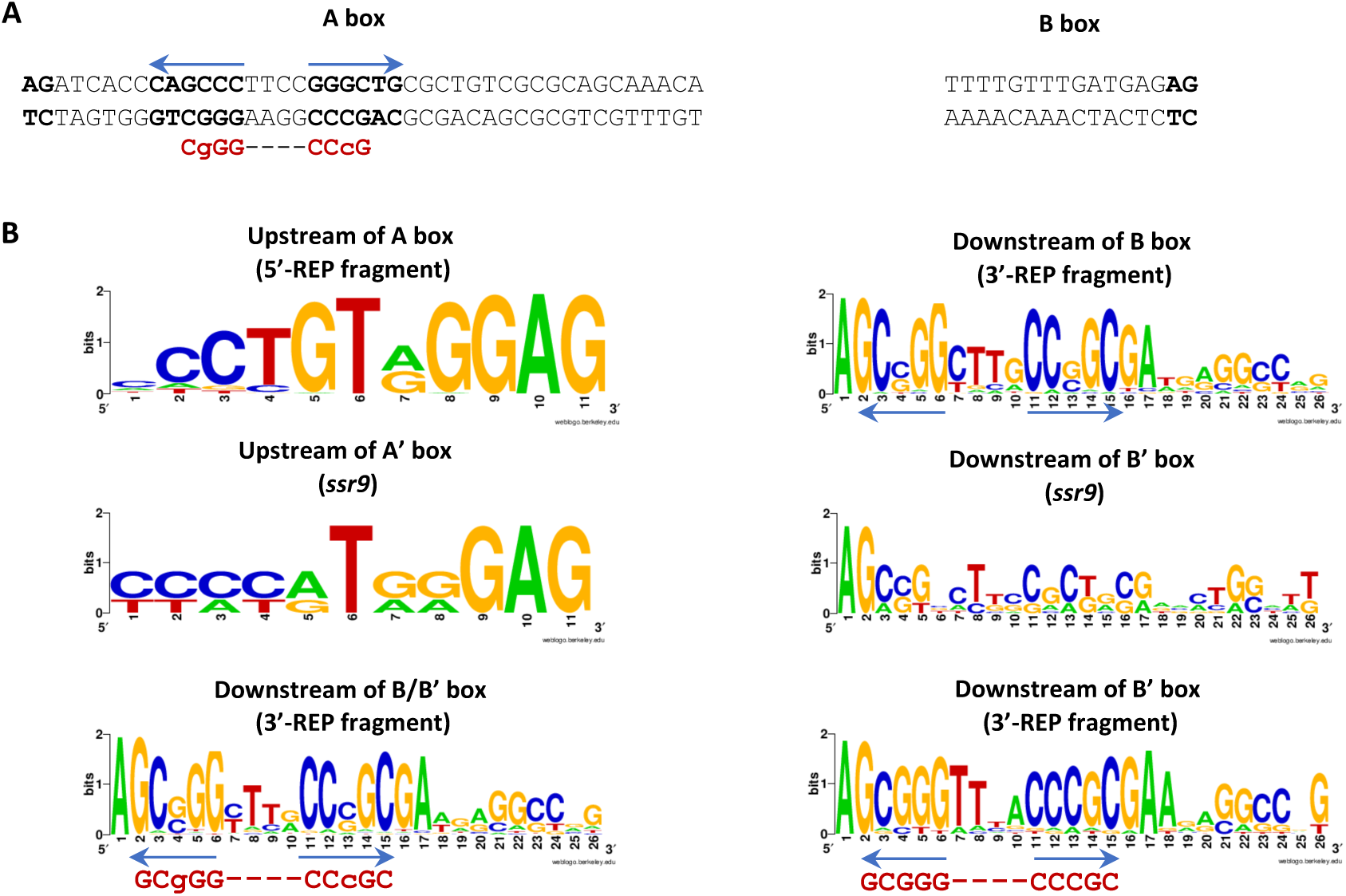
(**A**) Sequence of the ISPpu9 A and B boxes. The AG dinucleotide at the 5’-end of the A box and at the 3’-end of the B box are highlighted in bold. An inverted repeat present in the A box is shown with blue arrows and in boldface. (**B**) Sequence similarity upstream of A boxes (5’-REP fragment), or downstream of B and/or a B’ boxes (3’-REP fragment), at the insertion sites of the ISPpu9-like elements present in the bacterial strains listed in Tables S4 and S5. The similarities upstream of an A’ box, or downstream of a B’ box of the orphan *ssr9* genes present in *P. putida* strains KT2440, KBS0802 or NCTC13186 are also shown (indicated as “*ssr9*”). The short palindrome indicated in red downstream of the B or B’ boxes (3’-REP fragment) closely resembles that in the A box (Fig. 2A). Sequence comparisons and logos were generated with WebLogo (49) at http://weblogo.berkeley.edu.

This work shows that the A and B boxes are conserved in other closely related ISs. ISPpu9 was found to form several different circular intermediates that can explain the different combinations of the *asr9*, *tnp* and *ssr9* elements observed in KT2440 genome. Formation of these circular intermediates required the transposase, the terminal AG dinucleotides and short sequence patches at the IS ends and at the 5’-untranslated region (5’-UTR) of the *tnp* mRNA, which likely corresponds to the guide RNA. The possible formation of a new promoter upon joining of the IS end that might increase transcription of the *tnp* gene was analysed. The relevance of the Ssr9 sRNA for transposition, and its presence in other ISs closely related to ISPpu9, was also examined. The results presented provide insights into how the circular intermediates are formed and show that ISPpu9 differs in some aspects from the prototypical IS*110*, offering new perspectives on this peculiar family of ISs.

## MATERIAL AND METHODS

### Bacterial strains and culture media

The *Escherichia coli* strains DH5α (26), HB101 (pRK600) (27) and CC118λpir (28) were cultured in Lysogeny Broth (LB) (29) at 37°C. *P. putida* strains KT2440 and F1 (30,31) were cultured at 30°C in either LB or in M9 minimal salts medium (29) complemented with trace elements (32) and 30 mM citrate as the carbon source. Kanamycin (50 μg/ml), ampicillin (100 μg/ml), chloramphenicol (50 μg/ml) or streptomycin (50 μg/ml) were added as needed.

### Construction of delivery plasmids pKNG-ISPpu9-Km and pKNG-ISPpu9-Gm, transposition assays and identification of the insertion sites

A DNA fragment containing an ISPpu9 variant lacking *ssr9* (PP_4791), and with an XbaI site located between the end of the *tnp* gene and the start of the B box, and sites for NotI flanking both IS ends, was chemically synthesized and inserted into plasmid vector pUC57-mini (GenScript), obtaining plasmid pUC-ISPpu9ΔS. Determinants specifying for resistance to kanamycin or gentamycin were individually inserted at the XbaI site of pUC-ISPpu9ΔS, obtaining plasmids pUC-ISPpu9ΔS-Km and pUC-ISPpu9ΔS-Gm, respectively. These antibiotic-resistance genes were PCR-amplified from plasmids pSEVA225 (33) or pSEVA621 (33), respectively, using oligonucleotide pairs Km-D/Km-R or Gm/D/GmR (supplementary Table S1), which include restriction sites for XbaI. Plasmids pUC-ISPpu9ΔS-Km and pUC-ISPpu9ΔS-Gm were digested with NotI and the DNA fragments containing ISPpu9ΔS-Km or ISPpu9ΔS-Gm were individually cloned at the NotI site of the suicide delivery vector pKNG101 (34), obtaining plasmids pKNG-ISPpu9ΔS-Km and pKNG-ISPpu9ΔS-Gm. A similar delivery plasmid containing an ISPpu9 variant that includes the *ssr9* gene, named pKNG-ISPpu9-Km, has been described earlier (25).

The pKNG101-derived plasmids were introduced into *P. putida* F1 by conjugation in triparental filter mating assays, using plasmid pRK600 as the donor of transfer functions, as previously described (36). After 5 h at 30°C, the cells were suspended in 1 ml of M9 medium, and serial dilutions were plated onto agar plates containing M9 medium with citrate as the carbon source plus either kanamycin (for pKNG-ISPpu9ΔS-Km) or gentamycin (pKNG-ISPpu9ΔS-Gm). Since plasmid pKNG101 does not replicate in pseudomonads and bears a streptomycin resistance determinant, this marker was used to differentiate between Km-or Gm-resistant colonies in which the pKNG101-derivative had inserted into the chromosome by recombination from those in which ISPpu9 had transposed to the chromosome. Colonies being Km- or Gm-resistant and Sm-sensitive were presumed to derive from a true transposition event. The absence of pKNG101 in those strains was further monitored by PCR with oligonucleotides targeted to the *sacB* and Sm-resistance genes (Table S1). The presence of ISPpu9ΔS-Km or ISPpu9ΔS-Gm in the selected strains was checked by PCR using oligonucleotides targeted to the *tnp* gene, the Km-resistance or the Gm-resistance determinants (Table S1).

To identify the insertion site of ISPpu9ΔS-Km or ISPpu9ΔS-Gm in the chromosome of the F1-derivative strains obtained, total DNA was purified, partially digested with Sau3A-I, and the fragments obtained cloned at the BamHI site of plasmid pUC18 (35). The DNA insert was sequenced with oligonucleotides circ-5’-1 and 5’Km-seq, or circ-5’-1 and 5’Gm-seq (Table S1), which hybridize at the 5’ and 3’ ends, respectively, of the ISPpu9 *tnp* gene and of the Km- or Gm-resistance determinants, and allow sequencing outwards of these genes.

### Detection of ISPpu9- and ISPpu10-derived minicircles by PCR

Circular DNA intermediates were obtained from 4 ml of and O/N culture in LB medium, using the QIAprep Spin Miniprep kit (Qiagen). The DNA was eluted from the columns in 40 μl of 10 mM Tris (pH 8.5) and 1.5 μl were used as template in PCR reactions with the oligonucleotide primers indicated in the corresponding figures, which are oriented in opposite directions, facing outward the IS. Amplification was performed for 28 cycles, using the Promega PCR Mastermix. The amplified DNA fragments were visualized by electrophoresis in agarose genes, and later cloned into pGEM-T Easy (Promega) and sequenced. Plasmids containing the PCR-amplified DNA fragments corresponding to Junc-1 or Junc-2 minicircles were named pJunc-1 or pJunc-2, respectively.

### Real time qPCR

Total DNA was obtained from three independent cultures (biological replicas) of cells grown in LB medium to stationary phase, using the GNOME® DNA isolation kit (MP Biomedicals). Real-time PCR (qPCR) was performed using the 2^-ΔΔCt^ comparative method (36), as previously described (37,38), employing primers directed towards *rpoN*, *tnp*, *asr9* or *ssr9* genes, as specified (see Table S1). The results were normalized relative to those obtained for *rpoN*, which is present at one copy per genome.

### Transcriptional reporter fusions and ²-galactosidase activity assays

The transcriptional fusion of ISPpu9 promoter *Ptnp* to the *lacZ* reporter gene has been described earlier (25). The promoter regions of the ISPpu10 *tnp* and *asr10* genes were PCR-amplified with oligonucleotides Ptnp10dir and Ptnp10revH, or Pasr10dir and Pasr10revH, respectively, and cloned into plasmid pGEM-T Easy. The insert was excised as a EcoRI/HindIII fragment and cloned between the same sites of pSEVA225.

To analyse the transcriptional activity at the Junc-1 and Junc-2 minicircles, DNA fragments spanning the regions B-A’-*ssr9*-B’-A-*Ptnp* (Junc-1), or B-A-*Ptnp* (Junc-2), and whose 3’ end extended just up to -but not including-the *tnp* translational start site, were PCR-amplified using as a template plasmids pJunc-1 or pJunc-2, and the oligonucleotide pairs circ-3’-2/PtnpR or circ-3’tnp/PtnpR, respectively (see Supplementary Table S1). The amplified DNA segments were cloned between the BamHI and HindIII sites of plasmid vector pSEVA225 (39), which contains a promoter-less *lacZ* gene downstream of the HindIII site. The reporter plasmids obtained were named pPjunc1 and pPjunc2, respectively. An additional reporter plasmid containing only the “B-A” junction (lacking *asr9* and *ssr9*), was obtained amplifying a DNA fragment form a ISPpu9-derived minicircle with oligonucleotides BAjunc-rev and BAjunc-dir (Table S1); the amplified DNA fragment was cloned into pGEM-T Easy, excised from the resulting construct as an EcoRI segment, and cloned at the EcoRI site of pSEVA225, obtaining plasmid pSEVA-BA. The direction of the insert was verified by DNA sequencing.

To analyse the presence of a promoter at the junction region of the ISPpu10 minicircle, a DNA segment corresponding to the junction was PCR-amplified as indicated above (see section on minicircles), using oligonucleotides 5’-cir10 and 3’-cir10, and cloned into plasmid pGEM-T Easy, obtaining plasmid pJunc10. After sequencing to discard undesired mutations, the insert was excised as a EcoRI fragment and cloned at the EcoRI site of pSEVA225; obtaining pSEVA-Junc10. The direction was verified by DNA sequencing.

All plasmid constructs were sequenced to verify the absence of undesired mutations. The reporter plasmids were introduced into *P. putida* strain KT2440 as described earlier (25), and ²-galactosidase activity was assayed using *o*-nitrophenyl-²-D-galactoside (ONPG) as a substrate (40). A minimum of three independent assays (biological replicates) were performed.

### Generation of ISPpu9 mutant derivatives

Variants of ISPpu9 lacking the REP sequences upstream and downstream of the IS ends were obtained by PCR amplification of the ISPpu9 copy annotated as PP_4791, which lacks *ssr9*, and using appropriate oligonucleotides that introduced the desired mutations (Table S1). Additional variants with deletions or point mutations at the A box, the B box, the 5’-UTR region of *tnp* mRNA, or a mutant *tnp* gene (named *tnp*\*) that has three consecutive stops codons (TAATGATAA) after the 16^th^ codon, were chemically synthesized and inserted into plasmid vector pUC57-mini (GenScript). All these variants were also based on the ISPpu9 copy annotated as PP_4791, and include an engineered XbaI site located between the end of the *tnp* gene and the start of the B box, and sites for Not I at both ends of the DNA sequence. The corresponding DNA fragments were excised from their pUC57-mini vector with NotI and introduced at the NotI site of the suicide delivery plasmid pUT-Mini-Tn5Km (41). The resulting plasmids were introduced into *P. putida* F1 by conjugation using the helper plasmid RK600 (27) as donor of transfer functions. Since the donor plasmid does not replicate in *P. putida*, the kanamycin resistant transconjugants obtained should contain the ISPpu9 mutant variants inserted in the chromosome by way of the MiniTn5 transposon. Two clones of each variant were selected, and the presence of the Tn5 transposon and the mutations introduced were confirmed by PCR with appropriate test primers.

## RESULTS

### Conservation of the A/A’ and B/B’ boxes in ISs related to ISPpu9

As explained above, the *tnp* gene in all seven copies of ISPpu9 of strain KT2440 is flanked by sequences, named A and B boxes, that also flank the *ssr9* gene (designated A’ and B’ in this context) located downstream of the *tnp* gene. Interestingly, the three additional *ssr9* copies located at other chromosome loci, unlinked to ISPpu9, also carry these A’ and B’ boxes (25; Fig. 1 and Supplementary Table S2). While the seven A and B boxes were identical in sequence, the A’ and B’ boxes showed 1-3 mismatches relative to the A and B boxes (Table S2). In addition, alignment of the A/A’ boxes showed the presence of a conserved inverted repeat, a feature that is not evident at the B/B’ boxes (Fig 2A).

In all cases, an AG dinucleotide is present at the 5’ end of the A and A’ boxes, and at the 3’ end of the B and B’ boxes; this dinucleotide defines the boundaries of ISPpu9, or of the independent *ssr9* copies (shown in bold in Table S2). To evaluate the potential relevance of these sequences, their conservation in other ISPpu9-like elements was analyzed by performing a BLASTP search at the Pseudomonas Genome Database (42) using the ISPpu9 *tnp* gene copy annotated as PP_1133 as query. Ten *Pseudomonas* strains contained ISs with a transposase showing >87% amino acid identity to that encoded in PP_1133: *P. putida* KT2440 (7 copies), *P. putida* NCTC13186 (7 copies), *Pseudomonas sp.* KBS0802 (7 copies), *P. putida* B4 (3 copies), *P. putida* BIRD-1 (1 copy), *P. putida* S12 (4 copies), *P. putida* JB (1 copy), *P. plecoglossicida* XSDHY-P (19 copies), *P. putida* S16 (1 copy) and *P. putida* DLL-E4 (1 copy) (Table S3). In all cases, the transposase gene (*tnp*) was flanked by A and B boxes, with nucleotide identities of >87% (A box; Table S4) or 100% (B box; Table S5). Upstream of the *tnp* gene, a sequence showing a >95% identity to *asr9* was present in all cases. As reported earlier (25), the *ssr9* sRNA could only be found in strains KT2440, NCTC13186 and KBS0802, and was always flanked by A’/B’ boxes. Intriguingly, these three strains have seven copies each of ISPpu9, a number that is unlikely to be a coincidence. The gene context for each of these ISPpu9 copies was identical in the three strains (Fig. S1, supplementary material), suggesting that KT2440, NCTC13186 and KBS0802 derive from a common ancestor that already contained the seven ISPpu9 copies. Their genome sizes are similar, but not identical (6.18, 6.13 and 6.20 Mb, respectively (42)).

In a complementary approach, the presence of A or B boxes in *Pseudomonas sp*. genomes was analyzed using BLASTN at the Pseudomonas Genome Database (42), using the A and B boxes of the ISPpu9 copy annotated as PP_1133 as queries. Hits were retrieved for 21 strains. However, only in 10 strains the A/A’ boxes were associated to either an IS*110*-family *tnp* or to a *ssr9*-like gene, which were already identified when using the ISPpu9 transposase sequence as query (Table S4). In the case of *P. putida* KT2440, the search retrieved 16 matches. Seven corresponded to the known A boxes at the 5’ ends of the 7 ISPpu9 copies, and 8 to the A’ box upstream of the *ssr9* copies present in this strain. The last of these 16 matches corresponded to an incomplete A’ box (35 out of 43 bp) in a large intergenic region downstream of PP_5726 (*tki8*), which was not associated to a *tnp* or *ssr9* gene. Sixty-nine bp downstream of it, a B’ box was present, suggesting that this is a remnant of an isolated *ssr9* gene that suffered a deletion event. The remaining strains containing A/B boxes corresponded either to *P. putida*, closely related species such as *P. mosselii* or *P. plecoglossicida*, or unidentified *Pseudomonas* species (3 strains, one of which -KBS0802-is, as noted earlier, a very close relative of *P. putida* KT2440). A clear sequence conservation was observed in the A/A’ boxes detected, which included the imperfect inverted repeat shown in Fig. 2A.

A similar BLASTN search retrieved sequences identical to the B box in 19 strains, and in 17 of them the sequence was located downstream of an IS*110*-family *tnp*. However, in 9 of these 17 strains, all of which belonged to the *P. syringae* group, a clear A box upstream of the *tnp* gene was not detected, possibly due to mutations rendering it unrecognizable. These 9 strains were excluded from further analyses. Two additional *P. putida* strains, BIRD1 and JB, presented a DNA sequence adjacent to an IS*110*-family *tnp* that matched a B box but contained a 1 bp deletion compared to the query. Finally, B’-boxes were only found in the three strains containing *ssr9*, KT2440, KBS0802 and NCTC 13186 (Table S5). Interestingly, the isolated *ssr9* copy next to gene PP_4878 in strain KT2440 was followed by a small sequence corresponding to the end of an ISPpu9 *tnp* gene, which in turn ended with a B box. Therefore, the sequence was A’-*ssr9*-B’-‘*tnp*-B.

### Analysis of the target for ISPpu9 transposase

Since the seven ISPpu9 copies in KT2440 are inserted within a conserved REP sequence, it was hypothesized that ISPpu9 transposase exhibits high target specificity, inserting between the ninth and the tenth nucleotide of a REP sequence (10). To further extend this observation, the insertion site of 47 ISPpu9-like ISs listed in Table S3 was analyzed. Comparison of the nucleotide sequences flanking the insertion sites identified highly conserved positions that were strongly favoured (Fig. 2B). A palindromic sequence was evident in the REP sequence fragment downstream of the B or B’ boxes, as indicated by grey arrows in Fig. 2B (bottom panel). Notably, this palindrome exhibited strong similarity to that present within the A box (Fig. 2A).

To experimentally validate the observed insertion specificity, two suicide delivery plasmids were constructed. These plasmids carried the ISPpu9 copy annotated as PP_4791, which lacks *ssr9*, and had been engineered to include either a kanamycin resistance marker (plasmid pKNG-ISPpu9ΔS-Km), or a gentamicin resistance marker (plasmid pKNG-ISPpu9ΔS-Gm), immediately downstream of *tnp* and upstream of the B box. These plasmids were independently introduced into *P. putida* strain F1, which lacks ISPpu9 but contains over 300 intergenic REP sequences highly similar to those in *P. putida* KT2440. Isolates containing either ISPpu9ΔS-Km or ISPpu9ΔS-Gm, but lacking the delivery plasmid (which replicates in *E. coli*, but not in *Pseudomonas*), were obtained. The insertion sites were determined for three isolates containing the ISPpu9ΔS-Km element (named F1-21, F-22 and F1-36, Fig. 3A), and for two isolates containing ISPpu9ΔS-Gm (named F1-10 and F1-28; Fig. 3A). In all cases, the IS was inserted within a REP element located at an intergenic region. The insertion occurred at an AG dinucleotide immediately following position 9 of the REP element, with the IS boundaries including an AG at both the 5’-and 3’ ends. Reconstruction of the REP sequence at these insertion sites revealed high sequence conservation among the analyzed clones, although several mismatches were observed. Notably, the inverted repeat downstream of the AG dinucleotide, which serves as insertion site, was highly conserved (Fig. 3B, red font). Furthermore, the REP elements in these sites closely resembled the insertion sites for ISPpu9 in *P. putida* KT2440 (Fig. 3B; the consensus sequence for the 7 ISs of this strain is provided as reference). These findings offer strong experimental evidence that ISPpu9 has high target specificity for transposition.

**Figure 3.**
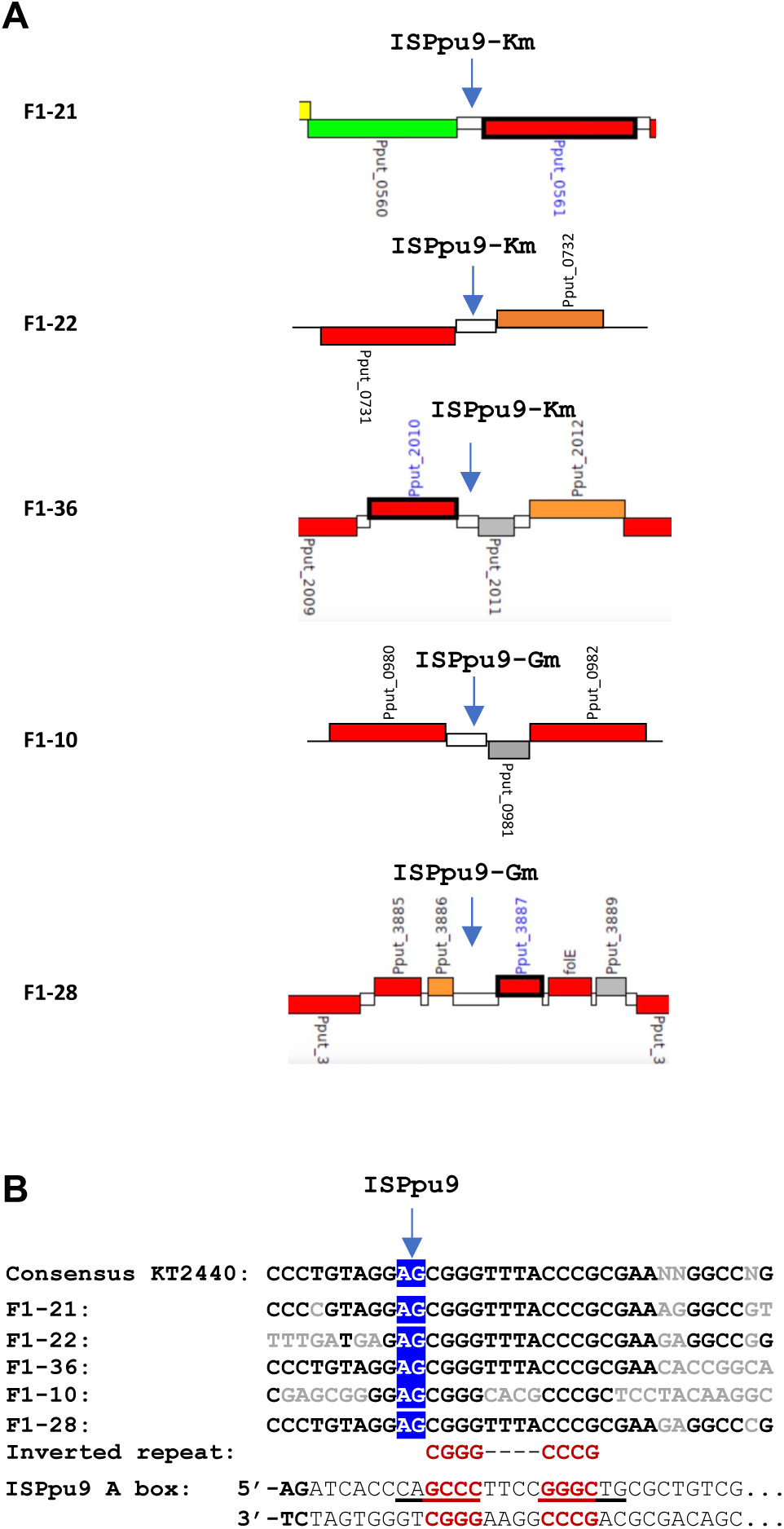
Identification of ISPpu9 insertion sites in five independent *P. putida* F1 isolates. The suicide delivery plasmids pKNG-ISPpu9ΔS-Km or pKNG-ISPpu9ΔS-Gm were individually conjugated into *P. putida* F1. Insertion sites were identified for three isolates with ISPpu9ΔS-Km, and two with ISPpu9ΔS-Gm, as described in Methods. (**A**) The blue arrow marks the IS insertion site, located in intergenic regions (white rectangle); flanking genes are shown in coloured rectangles. (**B**) Sequences of each insertion site highlighting in blue the AG dinucleotide at which the IS is inserted. The consensus REP sequence flanking the ISPpu9 copies in strain KT2440 is shown for comparison. Nucleotides differences between insertion sites are in grey.

### ISPpu9 forms several multiple circular intermediates in *P. putida* KT2440

The presence of ISPpu9 copies with or without a *ssr9* gene, along with the independent *ssr9* gene copies flanked by A/A’ and B/B’ boxes, suggested that these elements might transpose in a modular fashion. To investigate this possibility we leveraged the observation that several members of the IS*110* family can excise from their insertion sites, forming double-stranded DNA circular intermediates believed to act as transposition intermediates (15,16,20,21). To determine whether ISPpu9 can generate such circular intermediates, and to characterize their diversity, we designed oligonucleotide primers targeting various IS regions, with 3’ends facing outward. PCR amplification with these primers should yield a product only if the ends of ISPpu9 have joined to form a circular intermediate (primers indicated in Fig. 4A). The primers were also designed to test whether alternative circular intermediates could form (i) between the A and B’ sites (a minicircle containing asr9, *tnp*, and ssr9), (ii) between the A and B sites (a minicircle containing asr9 and *tnp*), or (iii) between the A’ and B’ sites (a minicircle including ssr9 alone). Genomic DNA from *P. putida* KT2440 (containing ISPpu9 and *ssr9*) and *P. putida* F1 (lacking ISPpu9 or *ssr9*; negative control) was used as PCR template. As shown in Fig. 4B, all primer pairs yielded amplification products with DNA from KT2440, but not from F1, as expected. The PCR fragments were cloned into the pGEM-T Easy plasmid, and inserts from 6-10 independent transformants per reaction were sequenced. The sequences confirmed the existence of several minicircles.

**Figure 4.**
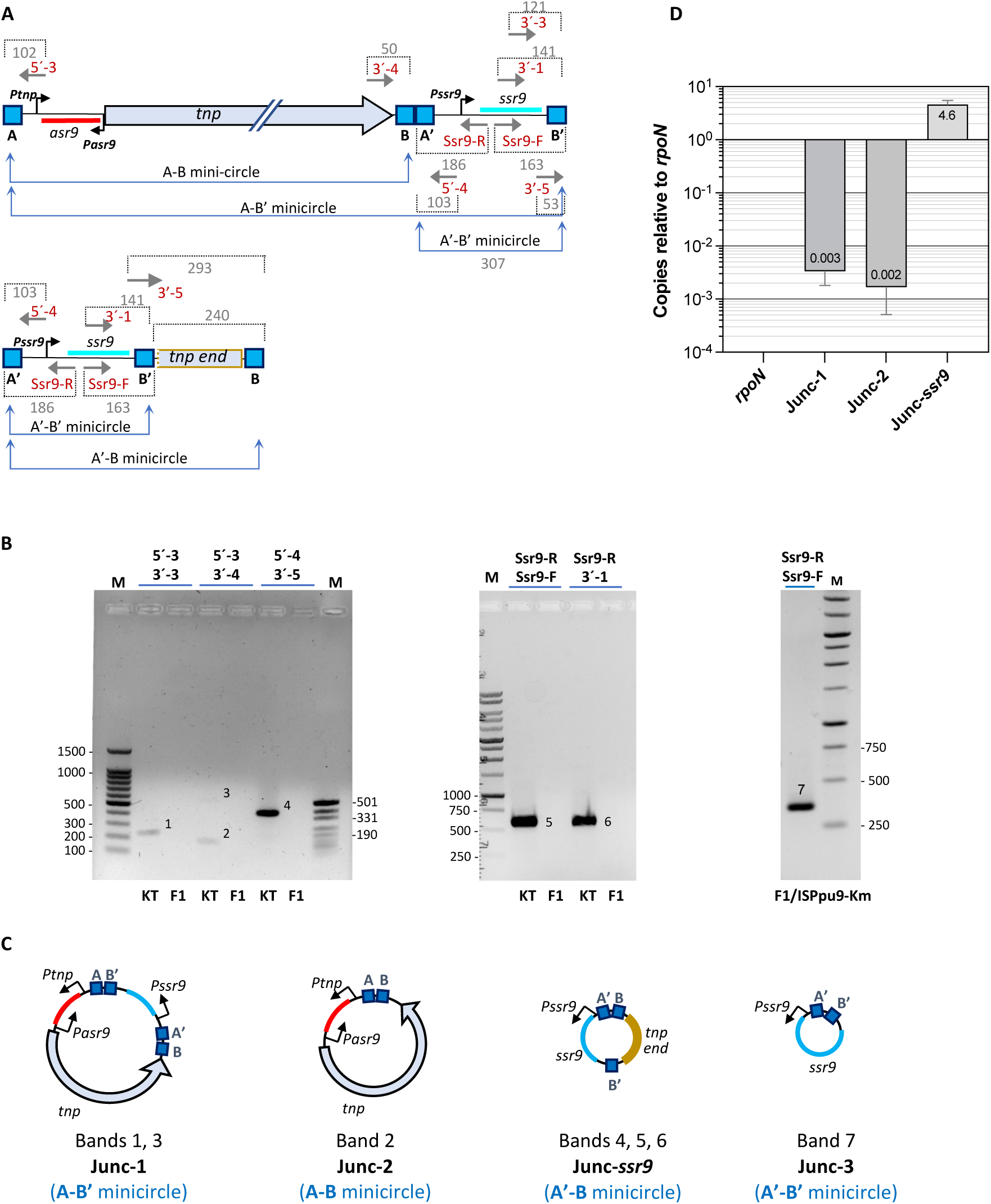
Identification of ISPpu9 circular intermediates. (**A**). Gray arrows represent primers (names in red font) used to PCR-amplify the predicted minicircles junctions formed by recombination of the A and B, A and B’, or A’ and B’ boxes. The distance, in base pairs, between the 5’-end of the primer and the 5’-/3’-end of the corresponding box, including the “AG” dinucleotide that flanks each box, is indicated (in the BA’ junction, the B and A’ boxes share the same AG dinucleotide). (**B**) PCR products obtained with the indicated primer pairs, resolved by agarose gel electrophoresis. PCR reactions were performed with DNA samples obtained from either strain KT2440 (indicated as “KT”), which contains ISPpu9 and *ssr9*, from strain F1 (lacks ISPpu9 or *ssr9*), or from an F1 isolate that has an ISPpu9-Km copy including *ssr9*, introduced by conjugation. (**C**) Minicircles interfered from the sequences of the different PCR-amplification products detected in (B). (**D**) Number of copies per genome of the Junc-1, Junc-2 and Junc-*ssr9* minicircles, determined by qPCR.

The largest minicircle, named Junc-1, resulted from recombination between the A and B’ boxes (containing *asr9*, *tnp* and *ssr9*), with a crossover at the AG dinucleotides located at the 5’-end of the A box and the 3’-end of B’ box (bands 1 and 3 in Fig. 4B; scheme in Fig. 4C). Another minicircle, Junc-2, formed by recombination between A and B boxes, contained only *asr9* and *tnp* (band 2 in Fig. 4B). A third minicircle, Junc-*ssr9*, resulted from recombination between an A’-box and a B box, likely derived from an independent *ssr9* copy located between genes PP_4877 and PP_4879, which is associated to a truncated *tnp* fragment (See Fig. 4A; bands 4, 5 and 6 in Fig. 4B). In this case, the amplified fragment included part of the 3’-end of *tnp*.

Interestingly, the PCR amplification product corresponding to Junc-*ssr9* was more abundant than that of Junc-1 and Junc-2. Quantitative PCR (qPCR) analysis, normalized against the *rpoN* gene (present at one copy per chromosome), revealed that Junc-*ssr9* was present at approximately 4.6 copies per cell, whereas Junc-1 and Junc-2 were ∼1800 times less abundant, occurring in only 1 out of 300–500 cells (Fig. 4D).

Unexpectedly, no minicircle corresponding to recombination between A’ and B’ boxes, flanking the independent *ssr9* copies in KT2440, was detected. This might be due to the high abundance of Junc-*ssr9*, which could mask detection of a less frequent A’-B’-type minicircle (hereafter Junc-3). To resolve this issue, an ISPpu9 variant containing *ssr9* was introduced into strain F1, which lacks independent *ssr9* copies, using the delivery plasmid pKNG-ISPpu9-Km. In this strain, a Junc-3 minicircle was detected, demonstrating that recombination between A’ and B’ ends can also occur (Fig. 4B, band 7).

### ISPpu9 circular intermediates in *P. putida* F1

Circular intermediates were also analyzed in three *P. putida* F1 derivatives containing ISPpu9 copies lacking *ssr9*. These included strains F1-21 and F1-36 (ISPpu9ΔS-Km) and F1-28 (ISPpu9ΔS-Gm) (see Fig. 3). Since these strains lack *ssr9*, only Junc-2-type intermediates (A-B box recombination) were expected. PCR assays confirmed the expected 366 bp amplification product for strains F1-21 and F1-36 (Fig. 5A–C). However, in F1-28, which was predicted to generate a 266 bp fragment, the observed fragment was unexpectedly larger (∼360 bp). Sequencing of the amplified DNAs showed that, in F1-21 and F1-36, minicircles resulted from the joining between the A and B boxes at the expected site. In strain F1-28, however, minicircles included an additional 150 bp sequence between the A and B boxes. Inspection of the ISPpu9ΔS-Gm insertion site in F1-28 indicated that excision occurred between the B box and a site 151 bp upstream of the A box, within a REP sequence identical to the one where ISPpu9ΔS-Gm had inserted (Fig. 5D and supplementary Fig. S2). The resulting minicircle contained an A box with its conserved AG dinucleotide, followed by 150 bp of chromosomal DNA that included a 5’-REP fragment at one end and a 3’-REP fragment at the other end, ending at the B box. This finding suggests that the ISPpu9 transposase may have mistakenly recognized the 3’-REP fragment located upstream of the A box with an identical sequence located 150 bp upstream. This might be a recurrent event since REP sequences are frequently organised as clusters of several elements with alternating orientations, known as BIMEs (Bacterial Interspersed Mosaic Elements)(43). This structural organization may increase the likelihood of misrecognition of the transposase, resulting in the formation of non-canonical circular intermediates.

**Figure 5.**
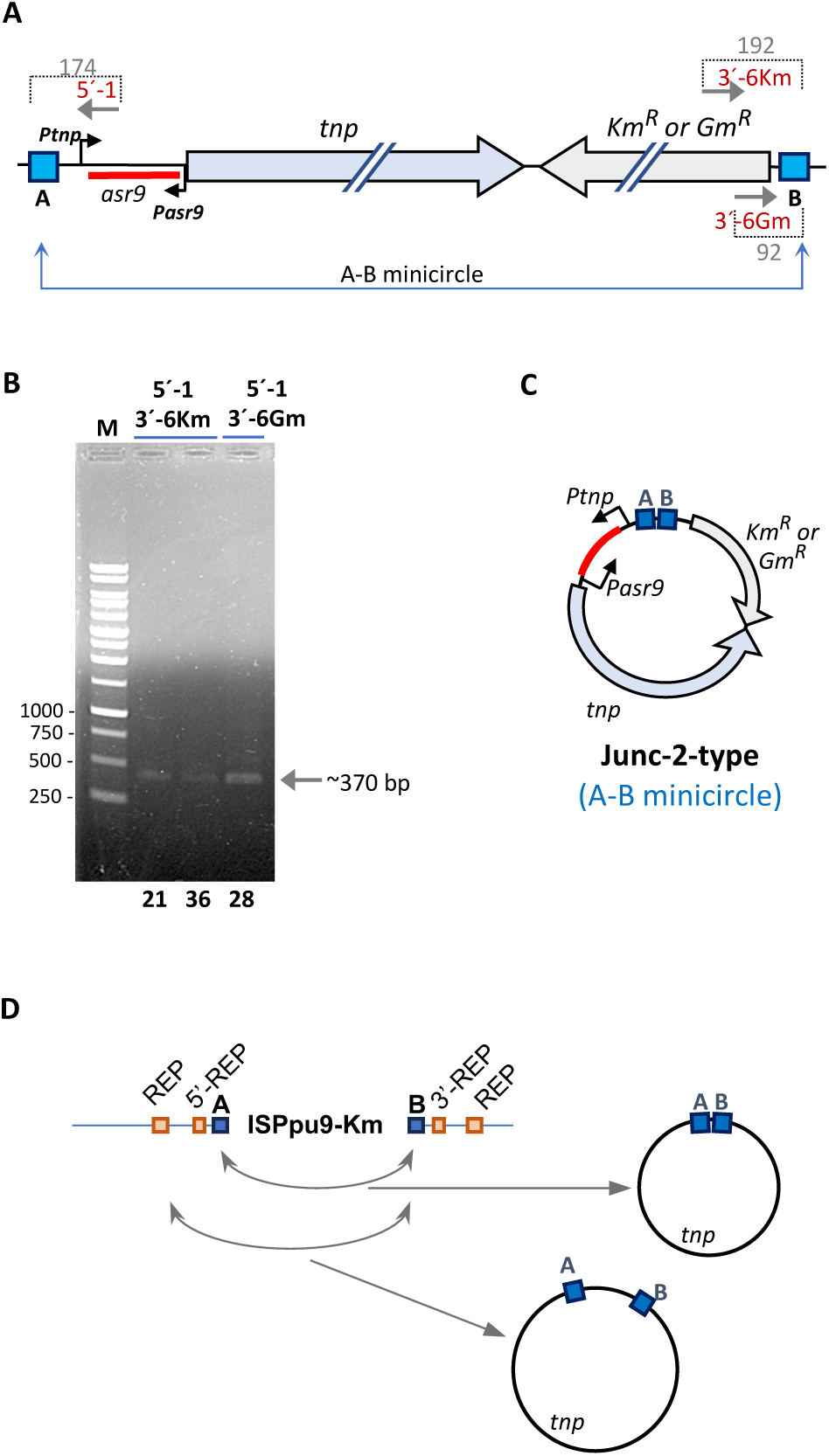
ISPpu9 circular intermediates in *P. putida* F1 derivatives containing ISPpu9. (**A**). Schemes of ISPpu9ΔS-Km and ISPpu9ΔS-Gm. Gray arrows show primers used for PCR amplification of predicted minicircle junctions formed by recombination of the A and B boxes. Numbers in grey indicate the distance in bp, between the 5’-end of the primer and the 5’/3’-end of the corresponding box, including the “AG” dinucleotide that flanks each box. (**B**) Minicircles detection in *P. putida* F1 derivatives containing ISPpu9-Km (strains F1-21 and F1-36) or ISPpu9-Gm (strain F1-28). The PCR products obtained with primer pairs 5’-1 and 3’-6km (for ISPpu9ΔS-Km), or 5’-1 and 3’-6Gm (for ISPpu9ΔS-Gm), were resolved by agarose gel electrophoresis. Lane M, DNA sequence ladder. (**C**) Minicircle deduced from sequencing the PCR-amplification products detected in (B), corresponding to Junc-2 type minicircles (A-B junction). Note that in the minicircle derived from strains F1-21 and F1-36, the A and B boxes are joint, but in that derived from strain F1-28, the A and B boxes are separated by a 151 bp segment (see text for details). (**D**) Scheme of minicircles formation through recombination between the AG dinucleotide flanking B an A boxes, or a REP element upstream of the A box identical to that at which ISPpu9 had inserted.

### The ISPpu9 circular intermediates do not generate a new promoter that increases *tnp* expression

As explained in the Introduction, the minicircles of several IS*110*-and IS*1111*-family ISs contain a new promoter, named *Pjunc*, generated through the joining of the right and left ends of the IS. The right end of the IS contributes the -35 region of the promoter while the left end provides the -10 region. This hybrid promoter increases transcription of the transposase gene (20–23). For IS*621* and IS*1111*, the hybrid promoter is believed to be the sole promoter driving transcription of the transposase gene in the minicircle (15,16). Among the minicircles identified in this study, Junc-1 and Junc-2 include the complete *tnp* gene. In Junc-1, *tnp* can be transcribed from promoters *Pssr9*, *Ptnp* and, if generated, a hypothetical new hybrid promoter formed by the joining of the B’ and the A boxes (Fig. 6A). In Junc-2, promoter *Pssr9* is absent, but *Ptnp* remains present (Fig. 6A).

**Figure 6.**
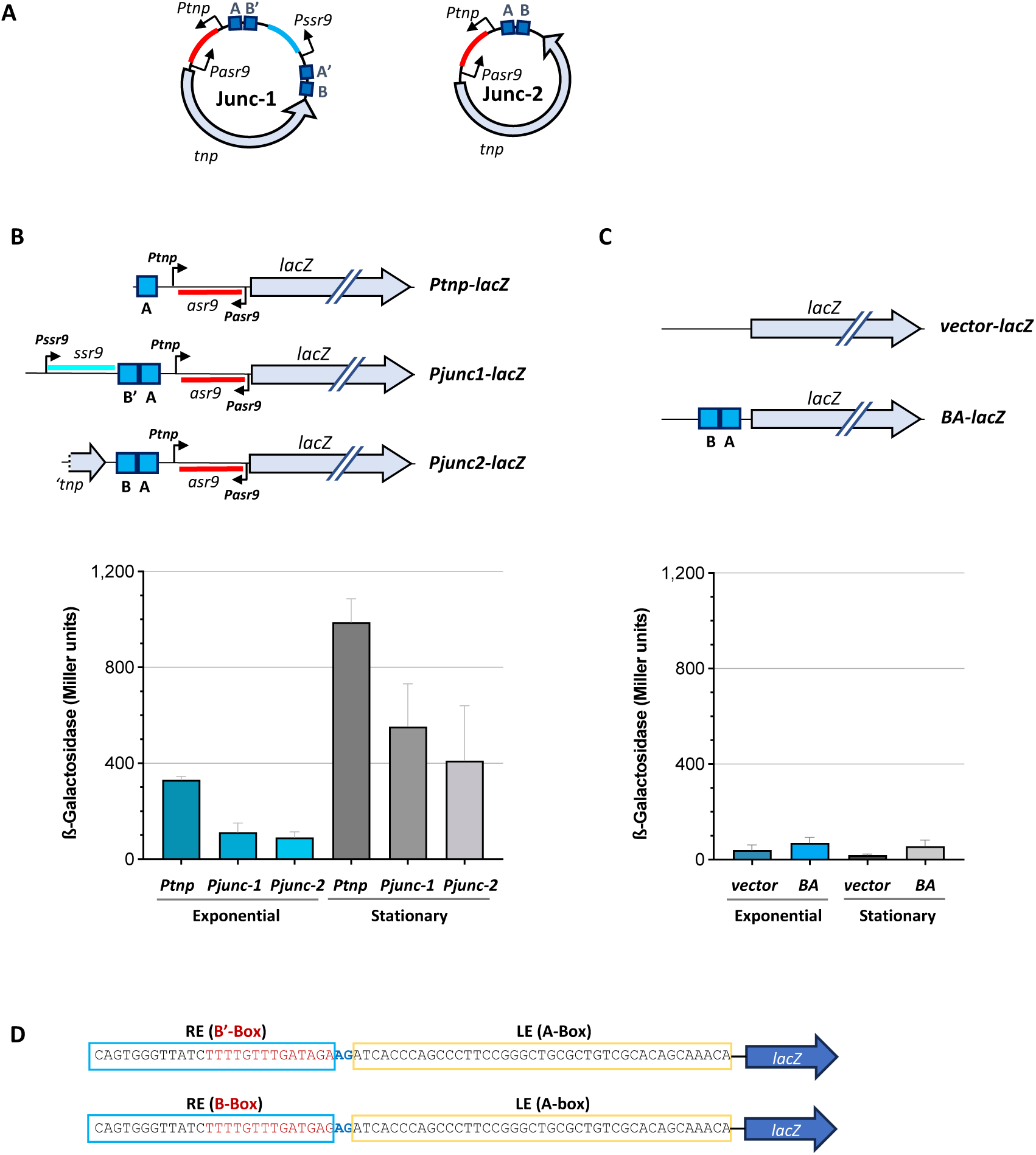
Transcription of the *tnp* gene in minicircles Junc-1 and Junc-2. (**A**) Schematics of Junc-1 and Junc-2 minicircles. (**B**) The transcriptional fusion *Ptnp-lacZ* has been described earlier (25), and includes the *Ptnp* promoter present in ISPpu9, fused to the *lacZ* reporter gene. Fusions *Pjunc1-lacZ* and *Pjunc2-lacZ* include the DNA regions located upstream of the *tnp* gene in the minicircles Junc-1 and Junc-2, respectively (the minicircles are depicted on the right side). The position of the *Ptnp* and *Pasr9* promoters, the genes specifying the *ssr9* and *asr9* sRNAs, as well as the A and B boxes, is indicated. The graph shows the ²-galactosidase activity of *P. putida* KT2440 cells containing plasmids pTNC-Ptnp (includes the *Ptnp-lacZ* fusion), pPjunc1 (includes the *Pjunc1-lacZ* fusion), or pPjunc2 (includes the *Pjunc2-lacZ* fusion), grown in LB medium to mid-exponential phase (A600 of 0.5), or early stationary (A600 of 2) phase. Error bars indicate standard deviations of three assays. (**C**) The transcriptional *BA-lacZ* fusion includes B and A boxes upstream of *lacZ*. All reporter plasmids derive from vector pSEVA225. (**D**) DNA sequences at B’-A and B-A junctions present in the reporter fusions shown in panels B and C. LE, left IS end (B and B’ boxes in red font); RE, right IS end.

To determine whether *tnp* transcription in any of these minicircles might be more efficient than in the chromosomal copies of ISPpu9, where *tnp* expression depends solely on the *Ptnp* promoter, two transcriptional fusions were constructed. These fusions, named *Pjunc-1-lacZ* and *Pjunc-2-lacZ*, contain the junction regions shown in Fig. 4 fused to the *lacZ* reporter gene in the orientation illustrated in Fig. 6A,B. The reporter plasmids were introduced into *P. putida* KT2440 and the ²-galactosidase activity was measured in parallel with a previously reported *Ptnp-lacZ* fusion, where *lacZ* expression depends exclusively on the native *Ptnp* promoter (Fig. 6B). In exponentially growing cells in LB medium, transcriptional activity from the promoter regions in junctions Junc-1 and Junc-2 was lower than that driven by the native *Ptnp* promoter (Fig. 6B). In stationary phase cells, ²-galactosidase activity increased approximately threefold in all reporter constructs. However, fusions containing *Pjunc-1* and *Pjunc-2* remained less active than the fusion containing only the native *Ptnp* promoter. These results suggest that the junctions formed by B’-A (Junc-1), or B-A (Junc-2), do not create a new hybrid promoter stronger than the native *Ptnp* promoter.

To further investigate this observation, the junction region containing only the B-A segment (lacking both *Pasr9* or *Ptnp* promoters), was cloned into the pSEVA225 reporter vector, just upstream of the *lacZ* reporter gene, generating the plasmid pBA-lacZ. As shown in Fig. 6C, ²-galactosidase activity was similar to that of the empty vector, confirming that no hybrid promoter is formed at the minicircle junction. Analysis of the DNA sequence of the B’A or B-A junctions (Fig. 6D) using the SAPPHIRE.CNN Pseudomonas-specific promoter prediction tool (44, available at https://sapphire.biw.kuleuven.be/index.php) did not predict the presence of a promoter.

It is worth noting that, in the Junc-1 minicircle, the transposase gene would be under the influence of both its native promoter (*Ptnp*) and the stronger *Pssr9* promoter (Fig, 6A) (25). However, this difference was not reflected in the activity of the *Pjunc-1-lacZ* reporter fusion, likely due to the presence of a transcriptional terminator at the end of *ssr9*, as predicted previously (25).

The absence of a hybrid promoter at the junction site contrasts with findings for IS*621*, the archetype of the IS*110* family, and with the observations of six other ISs in this family that generate hybrid promoters at their junction sites (15). This suggest that, ISPpu9 might be different from other members of the IS*110* family in this regard. To further explore this possibility, similar analyses were conducted for ISPpu10, another IS*110* family member present in *P. putida* KT2440. ISPpu10 encodes a transposase sharing 57.8% amino acid sequence similarity with ISPpu9 transposase. Additionally, ISPpu10 inserts at the same target site (a REP sequence) as ISPpu9 and has been shown to be active (10, 45). Interestingly, inspection of a previously published transcriptomic data for *P. putida* KT2440 (46) revealed abundant reads originating immediately upstream of the ISPpu10 transposase gene, and in the opposite direction, while the transposase gene presented very few reads (PP_1653; See Fig. 7A,B). This suggests that ISPpu10 specifies an antisense sRNA similar to Asr9 of ISPpu9, and that we therefore designated as Asr10.

**Figure 7.**
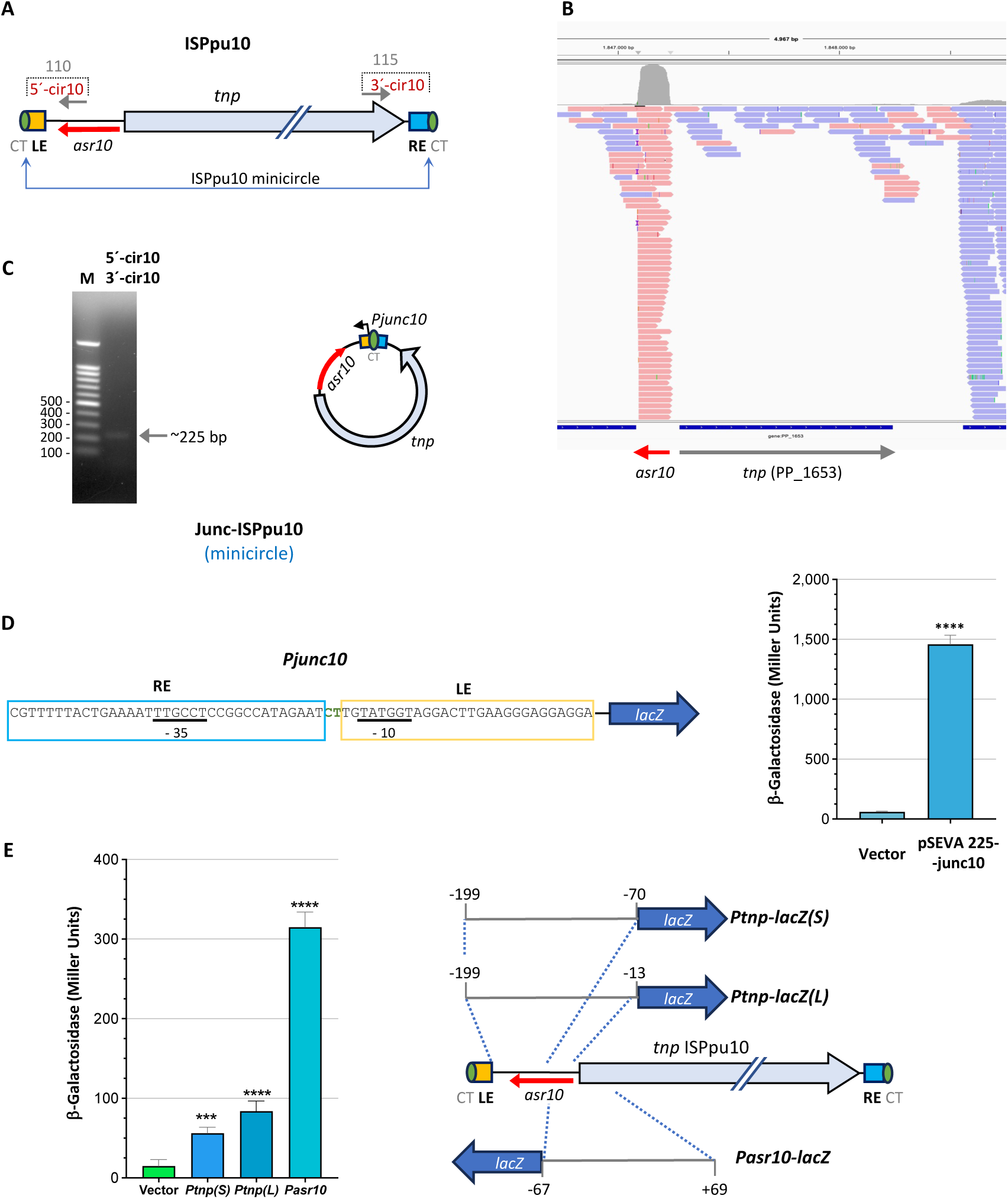
ISPpu10-derived minicircles. (**A**) Schematic representation of ISPpu10. The left end (LE) and right end (RE), as well as the CT terminal dinucleotide that defines the IS boundaries, are indicated. Outwards-facing oligonucleotides used to detect the minicircles formed by this IS and distances to IS ends are labelled. The *asr10* gene corresponds to the transcript reads detected in panel (**B**), which shows RNA-seq reads near the PP_1653 ISPu10 copy (pink: leftward transcription, blue rightward transcription). (**C**) DNA fragment detected after a PCR-amplification reaction using the primer pair indicated in panel (A) and a KT2440-derived lysate as substrate. The minicircle inferred after sequencing the amplified DNA fragment is depicted on the right. (**D**) ²-galactosidase assays in KT2440 cells growing exponentially (A_600_ of 0.6-0.8) in LB medium with either the empty reporter vector pSEVA225, or plasmid pSEVA225-junc10, containing the 225 bp PCR fragment amplified in panel (C) upstream of the *lacZ* gene, oriented as indicated on the left. Sequences derived from the right and left ends (RE, LE) are depicted, as well as the -35 and -10 consensus sequences for α^70^-RNA polymerase corresponding to a predicted hybrid promoter. (**E**) ²-galactosidase assays in KT2440 cells growing exponentially (A_600_ of 0.6-0.8) in LB medium and containing either the empty vector pSEVA225, or the reporter plasmids derived from pSEVA225 including the transcriptional fusions *Ptnp-lacZ(S)*, *Ptnp-lacZ(L)*, or *Pasr10-lacZ*. The DNA fragments cloned in each case are indicated on the left. Data represent means of six independent assays; the standard deviation is indicated. Asterisks indicate significant differences compared to the empty vector (one-way ANOVA; \*\*\*\**P*<0.0001; \*\*\**P*<0.001).

The presence of ISPpu10-derived minicircles in *P. putida* KT2440 was investigated using primers facing outward from the left and right IS ends. A PCR-amplified DNA fragment was detected, which was cloned and sequenced, confirming a Junc-2-type minicircle (Fig 7C). This DNA fragment was also cloned into pSEVA225 to generate a transcriptional fusion to *lacZ.* Introduction of the reporter plasmid into revealed that the junction region contains a strong promoter (Fig. 7D). Prediction of the junction sequence using the SAPPHIRE.CNN Pseudomonas-specific promoter prediction tool (44) identified a *Pseudomonas* σ^70^-RNA polymerase consensus promoter with clear -35 and -10 boxes (*P*-value of 0.000196). The -35 region derived from the IŚs right end, and the -10 region from the left end (Fig. 7D). Thus, the ISPpu10 minicircle harbours a strong hybrid promoter capable of driving transcription of the transposase gene, similar to IS*621*.

To determine whether ISPpu10 also contains a native (non-hybrid) promoter for the *tnp* gene, as seen in ISPpu9, two DNA segments of different length, both containing the region located upstream of the *tnp* gene, were cloned into pSEVA225, generating plasmids pSEVA225-Ptnp(S) and pSEVA225-Ptnp(L). In parallel, another DNA segment presumed to contain a promoter for *asr10*, which specifies the Asr10 sRNA, was also cloned into pSEVA225 upstream of the *lacZ* gene, generating pSEVA225-Pasr10. As presented in Fig 7E, strong transcriptional activity was detected for *Pasr10-lacZ* fusion, consistent with the abundant reads observed in the transcriptomic data (Fig. 7B). In contrast, transcriptional activity detected for the *Ptnp(S)-lacZ* and *Ptnp(L)-lacZ* fusions was low, albeit higher than that of the empty vector (Fig. 7E). These findings suggests that the ISPpu10 *tnp* gene can be transcribed from a weak native promoter upstream of *tnp*, but that its minicircle provides a much stronger hybrid promoter enhancing *tnp* expression, as described for other members of this family but different from what we observed for ISPpu9.

### Circle formation requires the transposase, and specific nucleotide sequences in the A and B boxes

The strong conservation of the A and B boxes among the ISPpu9-related ISs analyzed (Tables S4 and S5), and the presence of the conserved inverted repeat (GCCCnnnnGGGC) both at the A box and at the 3’-REP fragment downstream of the B box (Figs. 2 and 3), suggested that these elements might be important for minicircle formation. To investigate this possibility, variants of ISPpu9 were constructed lacking the A box (ISPpu9ΔA), the B box (ISPpu9ΔB), the 5’-REP fragment (ISPpu9Δ5’REP), or the 3’-REP fragment (ISPpu9Δ3’REP). The AG dinucleotide that defines the IS ends was conserved in ISPpu9ΔB and ISPpu9Δ5’REP. In ISPpu9ΔA, the terminal AG dinucleotide was deleted, but the remaining sequence at the resulting 5’-end was TG<u>AG</u>GT, which includes an AG dinucleotide (underlined). The last variant, ISPpu9Δ3’REP, lacked the G of the AG dinucleotide, although it conserved the A, which was preceded by another AG. These variants are depicted in Figs. 8A,B. To test whether minicircles are generated by the transposase, an additional variant was made that included three consecutive stop codons after the 16^th^ codon of the *tnp* ORF (ISPpu9*tnp*\*). Since at least some these IS variants were suspected to be unable to transpose into *P. putida* chromosome, they were introduced into strain F1 chromosome using a mini-Tn5 transposon. To this end, the different variants, as well as a wild-type ISPpu9, were cloned into the suicide plasmid pUTmini-Tn5Km, which does not replicate in *Pseudomonas*. The resulting plasmids were used to deliver the mini-Tn5Km-ISPpu9 constructions to *P. putida* F1 chromosome. Several Km-resistant clones were obtained in each case, and the presence of the appropriate mini-Tn5Km-ISPpu9 variant was confirmed by PCR. Two independent clones of each IS variant were selected, and the presence of minicircles analyzed by PCR using primers C3 and C4, which were expected to render a 494 bp DNA fragment derived from a Junc-2-type minicircle.

**Figure 8.**
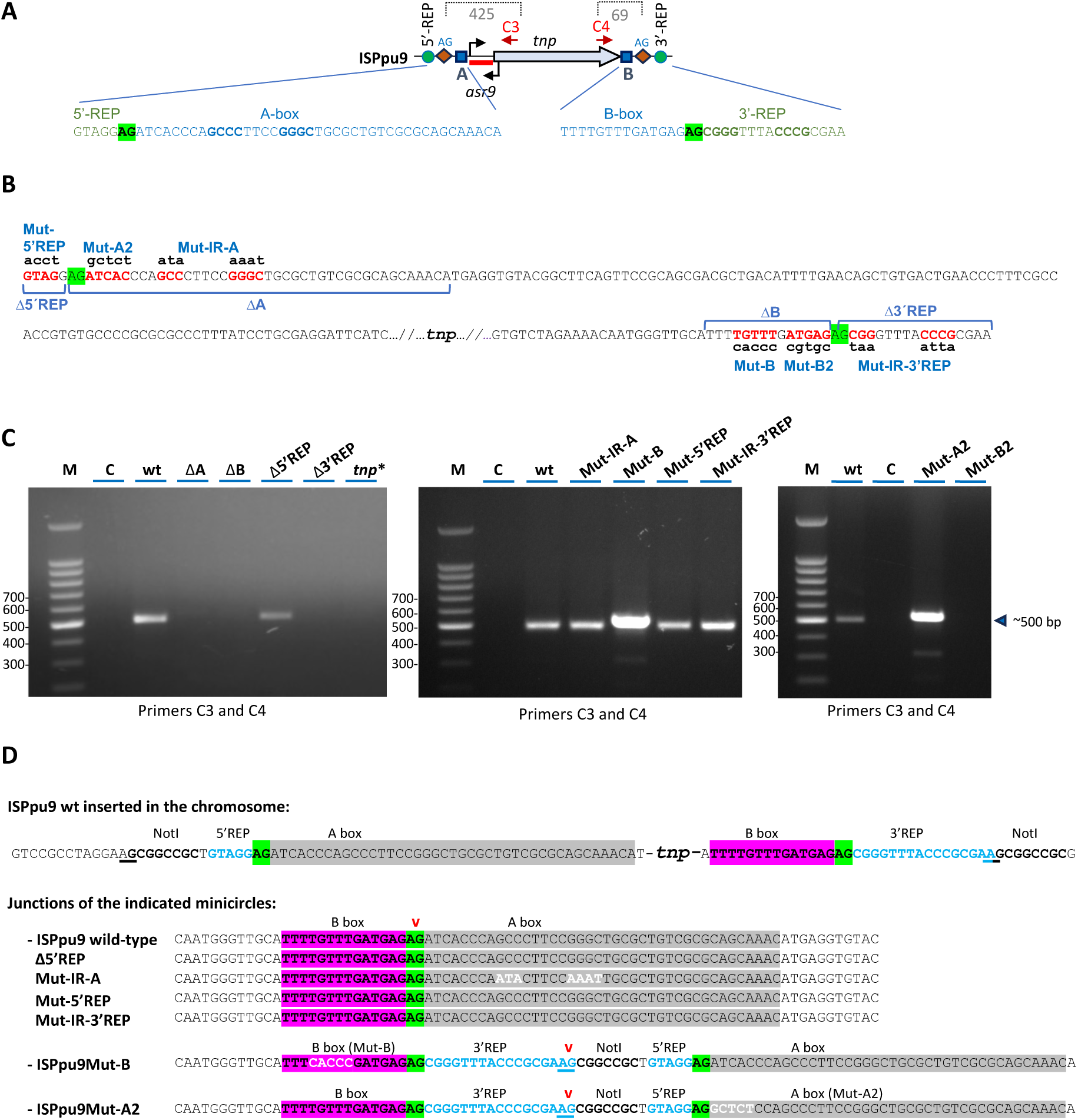
Influence of A box, B box, the 5’-and 3’-REP sequences, and the transposase on minicircles formation. (**A**) Scheme of ISPpu9 and the primers (C3 and C4) used for PCR detection of minicircle junctions. The distance between the 5’-end of the primers used and the end of the IS, including the flanking “AG” dinucleotides, is indicated. (**B**) Deletions or mutations introduced in ISPpu9 to obtain variants lacking A box (ISPpu9ΔA), B box (ISPpu9ΔB), the 5’-and 3’-REP sequences (ISPpu9Δ5’REP and ISPpu9Δ3’REP), or containing mutations in either A box (ISPpu9Mut-A2 and ISPpu9Mut-IR-A), B box (ISPpu9Mut-B1 and ISPpu9Mut-B2), or the 5’-or 3’-REP sequences (ISPpu9Mut-5’REP and ISPpu9Mut-IR-3’REP). ISPpu9*tnp\** contains three consecutive stop codons at the start of the *tnp* gene. (**C**) PCR amplification assay to detect minicircles in *P. putida* F1 strains containing either the wild type ISPpu9 (wt), or the ISPpu9 variants indicted in panel B, using the C3 and C4 primers indicated in panel A. M, DNA size ladder; C, control (no DNA added). The size of the amplified DNA fragments is indicated. (**D**) Sequence of the junction of the minicircles formed by the wild type ISPpu9 and the mutant variants. The sequence of the left and right ends of ISPpu9 when inserted into a REP sequence is indicated on top, flanked by artificially introduced sites for the NotI restriction enzyme for cloning purposes. The 5’ and 3’ ends of the REP sequence are in blue, A box in grey, B box in magenta; the underlined AG dinucleotides correspond to those believed to have been used in the recombination that led to the minicircle derived from ISPpu9Mut-B, ISPpu9Mut-A and ISPpu9Mut-B2.

The results are presented in Fig. 8C for one clone of each variant, as the result was consistent for both. Amplification bands of the expected size were obtained for the wild type ISPpu9 and the variant mutant ISPpu9Δ5’REP, and their sequence corresponded to a Junc-2-type minicircle in which recombination had joined the A and B boxes at the AG dinucleotide. No amplification products were detected when either the A box, the B box or 3’-REP element (plus the G of the terminal AG dinucleotide) had been deleted, suggesting that these DNA segments include nucleotides needed for the minicircles to form. Furthermore, the lack of transposase abolished the formation of minicircles.

To further delimit the sequences needed for minicircle formation, point mutations were introduced either at the inverted repeat present within the A box (ISPpu9Mut-IR-A), at the sequences immediately downstream of the AG dinucleotide of the left end of the IS (ISPpu9Mut-A2), or at specific positions of the B box (ISPpu9Mut-B and ISPpu9Mut-B2). In addition, to analyse the relevance of the 5’-REP and 3’-REP flanking sequences, mutations were introduced at the 5’-REP fragment (ISPpu9Mut-5’REP), or at the inverted repeat present in the 3’ REP fragment (ISPpu9Mut-IR-3’REP). These modifications, all of which conserve the flanking AG dinucleotide at both ends, are indicated in Figs. 8A,B. These variants were introduced into the *P. putida* F1 chromosome via the mini-Tn5 delivery transposon, and the minicircles formed were analysed by PCR as indicated above. Amplification products obtained for mutants ISPpu9Mut-IR-A, ISPpu9Mut-5’REP and ISPpu9Mut-IR-3’REP had the expected size and sequence, identical to that of the wild type (Fig. 8C,D). However, the amplified fragment derived from mutant variants ISPpu9Mut-B and ISPpu9Mut-A2 was somewhat larger. Sequencing of this DNA fragment indicated that the minicircle had arisen from the joining of sequences outside the IS boundaries. Specifically, the junction region included the mutant B box, followed by the terminal AG dinucleotide and 32 additional nucleotides containing the 3’-REP fragment, an AG dinucleotide, a target for the Not I restriction endonuclease, the 5’-REP sequence and the AG dinucleotide that corresponds to the A-box. This suggests that recombination had taken place between the AG dinucleotide present at the 5’-end of the two NotI sites artificially introduced in our constructs for cloning purposes, one located upstream of the 5’-REP fragment and the other one after the 3’-REP fragment (see Fig. 8D, the two AG used for recombination are underlined). This result was observed in several independent PCR amplifications and was, therefore, reproducible. Recombination appeared to be mediated by the transposase, rather than by a homologous recombination event between the two NotI sites, since these NotI sites were also present at ISPpu9 variants that were completely inactive (ISPpu9ΔA, ISPpu9ΔB and ISPpu9Δ3’REP; Fig. 8C).

The result suggests that the mutations introduced in mutant variants ISPpu9Mut-B and ISPpu9Mut-A2 do prevent the transposase from properly recognise the IS ends, but do not impair circularization. Instead, these mutations force the transposase to recombine at different but closely located sequences. However, what could facilitate a recombination at this alternative site is not obvious. Finally, no amplification product was obtained for mutant ISPpu9Mut-B2 (Fig. 8C), suggesting that the nucleotides of the right side of the B-box, which are adjacent to the AG dinucleotide that signals the 3’-end of the IS, are essential for circularization.

### Altering short sequences at the 5’-UTR of *tnp* mRNA complementary to IS ends impairs circularization

In the ISs of the IS*110*/IS*1111* family, the bridge RNA associated to the transposase contains two internal loops that base-pair with the nucleotides in the target DNA and two more that interact with the minicircle acting as donor, mediating recognition and recombination (15,16,24). In ISPpu9, the bridge RNA is predicted to be part of the non-coding region of the *tnp* mRNA. Analysis of the *tnp* 5’-UTR revealed three hexanucleotides sequences (marked with red spots in Fig. 9A) that are complementary to the short patches of the IS ends identified as critical for minicircle formation. Moreover, an inverted repeat highly similar to that present at the left end of the A-box and on the 3’REP was also evident (Fig. 9A, green spot). To investigate their roles, ISPpu9 variants in which these sequences were altered, were constructed (depicted in Fig. 9B). The variants targeting the IS ends were named ISPpu9AbB1, ISPpu9AbB2 and ISPpu9BbB, while the variant affecting the inverted repeat was named ISPpu9RB. These constructs were introduced into the F1 strain chromosome using the mini-Tn5 transposon described above, and their effects on minicircle formation was analysed by PCR. As shown in Fig. 9C, mutations in the bridge RNA sequences targeting the IS ends impaired minicircle formation, whereas mutations affecting the inverted repeat did not.

**Figure 9.**
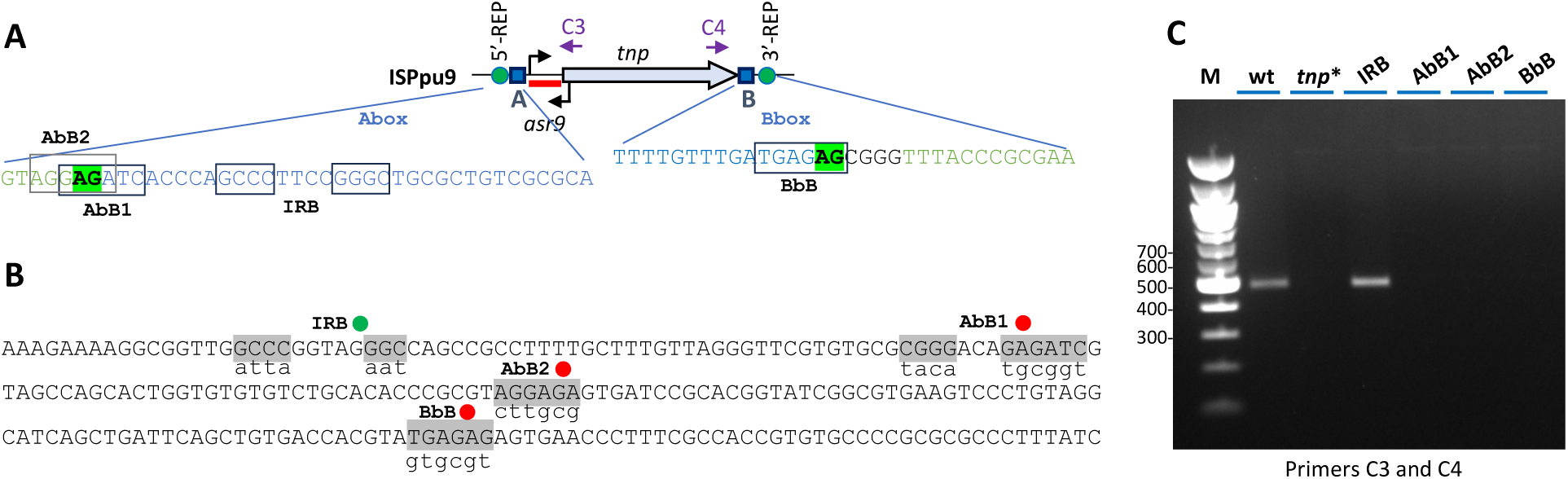
Influence on minicircle formation of sequences within the 5’-UTR of *tnp* mRNA that match sequences at the IS ends. (**A**) Schematic of ISPpu9, indicating the sequence stretches at the IS ends that are also found within the 5’-UTR of the *tnp* mRNA. These sequences, which are boxed, are named AbB1, AbB2, BbB and IRB. (**B**) Mutations introduced into the 5’-UTR of *tnp* mRNA, at the sites that match the AbB1, AbB2, BbB and IRB boxes. The original sequence is in uppercase, the altered one in lowercase. (**C**) PCR amplification assay to detect minicircles in *P. putida* F1 strains containing the wild type ISPpu9 (wt), or the ISPpu9 variants with mutations at the sequence boxes AbB1, AbB2, BbB, or at the IRB inverted repeat. An F1 strain containing ISPpu9*tnp*,* which cannot express the transposase, was used as negative control (*tnp*\*). Primers C3 and C4 were used for PCR detection of minicircle junctions, as indicated in Fig. 8.

## DISCUSSION

A notable feature of ISPpu9 is the presence of two variants in strain KT2440, differing by the presence or absence of the *ssr9* gene, as well as the presence of three isolated *ssr9* copies. Our results indicate that ISs elements closely related to ISPpu9 (transposase with >87% amino acid identity) are present in several *Pseudomonas* strains, although the *ssr9* gene was found only in *P. putida* strains KT2440, KBS0802 and NCTC13186. This suggests that *ssr9* is not essential for ISPpu9 functionality, an idea further supported by the successful transposition of an ISPpu9 variant lacking *ssr9* into the chromosome of *P. putida* strain F1, which lacks prior copies of either ISPpu9 or *ssr9*, but contains REP elements similar to those of strain KT2440 that can serve as targets. Despite the genomic differences among strains KT2440, KBS0802 and NCTC13186, all share 7 ISPpu9 copies inserted at intergenic positions within the same genomic context. The three isolated *ssr9* copies in strain KT2440 also appear in the same genetic context in strains KBS0802 and NCTC13186. This strongly suggests that these three strains derive from a common ancestor already harbouring the seven ISPpu9 copies and *ssr9*. Interestingly, some *ssr9* copies in strains KBS0802 and NCTC13186 are duplicated and appear in tandem (Fig. S2), raising the possibility that *ssr9* may move independently leaving the *tnp* gene behind. The detection of Junc-3-type minicircles containing just the *ssr9* gene supports this hypothesis.

The observation that all seven ISPpu9 copies in KT2440 are inserted at different chromosomal sites but consistently within intergenic REP sequences, initially suggested strong target specificity (10), although this was not experimentally tested at the time. Here, we used a suicide delivery vector to transfer ISPpu9 into *P. putida* F1, which lacks prior copies of this IS. Analysis of five insertion derivatives confirmed that ISPpu9 had integrated into intergenic REP elements with sequences highly similar to those of strain KT2440, demonstrating its target specificity. Some sequence variability was observed at the insertion sites in *P. putida* F1 target site, however. Other IS*110* family elements, such as IS*492* from *P. atlantica* (17,18) and IS*621* from *E. coli* (15,19), also integrate at specific sites, and some target flexibility has been noted. The target preferences differ for each IS*110*-family ISs, since target specificity is mediated by a bridge RNA that binds to the transposase, and interacts directly with the target DNA (15,16,24). Variations in the nucleotides on the recognition loops of the bridge RNA allow distinct target preferences, enabling each IS to recognize different, yet specific, targets.

REP sequences often form clusters, referred to as REPINs, which consist of REP pairs in opposite orientations separated by <51 bp, forming palindromic structures that can adopt stem-loop conformations (12,47). It has been proposed that REPINs may act as attractive targets for IS insertion (10,48). At least five of the seven ISPpu9 or ISPpu10 copies are inserted into a REPIN. Similarly, two out of three *P. putida* F1 derivatives containing the ISPpu9-Km were inserted into REPINs. This suggests that ISPpu9, like other IS*110*-family elements targeting REP sequences, may have a preference for REPINs in addition to sequence specificity.

A distinctive feature of ISPpu9 that sheds some light on its transposition mechanism is the presence of conserved flanking sequences, A and B boxes, at its ends, as well as analogous sequences (A’ and B’ boxes) flanking the *ssr9* gene. The 5’-end of the A/A’ boxes and the 3’-end of the B/B’ boxes are marked by an invariable AG dinucleotide. The results presented here show that ISPpu9 forms circular intermediates by joining its left and right ends, consistent with what has been found for other IS*110*/IS*1111* family members (15,16,20,21,22,23). However, four distinct minicircles were identified in strain KT2440, arising from recombination between various combinations of A, B, A’, and B’ boxes (Fig. 4). These minicircles include Junc-1 (A-B’ junction), Junc-2 (A-B junction), Junc-3 (A’-B’ junction), and Junc-*ssr9* (A’-B junction). This later minicircle was ∼1800 fold more abundant than Junc-1 or Junc-2 for no obvious reasons. The formation of these circular intermediaries explains the variety of ISPpu9-derived forms found on the KT2440 chromosome. The *ssr9* copy linked to a truncated fragment of the *tnp* gene might have arisen from a deletion within the Junc-1 minicircle including the A box, *asr9* and part of the *tnp* gene, which would generate a minicircle similar to Junc-*ssr9.* If such minicircle integrates at a chromosomal REP site, it would lead to a DNA segment comprising A’-*ssr9*-B’-‘*tnp*_end-B, identical to that found in strain KT2440 chromosome (see Fig. 4).

The formation of circular intermediates derived from IS elements often generates a new promoter, named *Pjunc*, by combining sequences from opposite ends that provide the -35 and -10 elements for recognition by RNA polymerase (20–23). This promoter can enhance transposase transcription, increasing its expression, and allows for the transcription of the bridge RNA that binds to the transposase and guides it to its specific DNA target (15, 16). In ISPpu9, however, our results show that *tnp* transcription does not increase in the Junc-1 or Junc-2 minicircles compared to the linear IS. The native *Ptnp* promoter remains active in these intermediates and is not weak. In fact, transposase expression appears to be post-transcriptionally regulated via its long structured 5’-UTR (25). In contrast, ISPpu10 has a weak native promoter and generates a strong hybrid *Pjunc* promoter that largely compensates the weakness of the native promoter. Thus, we propose that hybrid promoters are required only when native promoters are weak. The native promoter may provide sufficient transposase for initial minicircle formation, while hybrid promoters increase transposase expression in the minicircles to promote transposition.

Both RNA-seq assays and the activity of promoter *Pasr10* show that ISPpu10, as previously described for ISPpu9, generates an antisense sRNAs (Asr10). Inspection of ISPpu9-related ISs with >87% transposase identity suggests that similar antisense sRNAs may be widespread. Given its location, this sRNA may be involved not only in regulation of translation, as described for ISPpu9 (25), but perhaps also in the processing of the long RNA generated from promoter *Ptnp* that includes the bridge sRNA followed by the *tnp* mRNA, and that must be excised to generate the bridge RNA, as described for other IS*110* ISs (15,16).

An ISPpu9 derivative unable to express the transposase could not form minicircles, indicting that these are the product of a transposase-driven excision reaction. Eliminating or mutating the 5’-REP fragment, or modifying the 3’-REP fragment, still permitted minicircle formation if the terminal AG dinucleotides were conserved. This suggests that the target sequences recognized for insertion are not necessary for the excision step, during which a circular intermediate is formed via recombination of both ends of the IS, resulting in the loss of the target sequences.

ISPpu9 constructs lacking the A box or the B box were unable to form minicircles in *P. putida* F1, indicating that some specific nucleotides within these elements are required for the excision reaction. Point mutations at both ends of the IS either abolished minicircle formation or led to minicircles in which recombination had occurred at an unexpected site, utilizing AG dinucleotides at artificial NotI sites introduced during cloning. Finally, short sequence patches within the 5’-UTR of the *tnp* mRNA matched the sequence of the IS ends defined as critical for minicircle formation. This 5’-UTR includes the bridge RNA that should be processed and incorporated into the transposase to guide it to the target DNA. According to the current model for how the guide RNA recognizes the target, this RNA should have short sequences complementary to the top and bottom strands of the target to guide recognition (15,16,24). Mutating these short patches at the 5’-UTR of the ISPpu9 mRNA impaired minicircle formation, which supports the importance of the IS ends for minicircle formation. Characterization of the nucleotides required for minicircle formation, and the possible formation of non-canonical transposition intermediates through the use of closely related sequences, needs to be further analyzed, especially in the context of potential uses of these ISs in genome editing.

An intriguing observation is the presence of a short-conserved palindrome (GCCC-N_4_-GGGC) at three sites: the right target end (the 3’-REP fragment), within the ISPpu9 A box (close to the IS left end) and at the 5’-UTR region corresponding to the bridge RNA. It is tempting to speculate that this palindrome might play a role at some point in the transposition process. However, minicircles were still detected when this palindrome was altered by mutation at any of these three sites, and the junction sequence showed that recombination had occurred at the same site as in the wild type ISPpu9. While the palindrome may not be strictly required, it remains possible that it facilitates assembly of the recombination complex potentially enhancing excision efficiency. Alternatively, it may be dispensable for excision but necessary for integration.

Overall, the results presented show that ISPpu9 displays features characteristic of other IS*110*-family members, including target specificity or formation of circular intermediates. However, it also has distinctive features, such as the strong native promoter driving expression of the bridge RNA and the transposase mRNA, the ability to form multiple circular intermediates that enable modular transposition, and the presence of the Asr9 and Ssr9 sRNAs.

## DATA AVAILABILITY

The data underlying this article are available in the article and in its online supplementary material.

## SUPPLEMENTARY DATA

Supplementary data are available online (Max 2MB each file).

## AUTHOR CONTRIBUTIONS

Elena Parés: Methodology, Conceptualization, Formal analysis, Validation.

Luis Yuste: Methodology, Formal analysis.

Fernando Rojo: Conceptualization, Formal analysis, Visualization, Writing, Funding acquisition.

Renata Moreno: Conceptualization, Formal analysis, Visualization, Writing.

## ACKNOWLEDGEMENTS

We are grateful to J.C. Oliveros (CNB Bioinformatics Service) for excellent bioinformatic support.

