## Supplementary material for "The ISPpu9 insertion sequence of *Pseudomonas putida* KT2440 generates various circular intermediates enabling modular transposition"

**Supplementary information for:**

### The ISPpu9 insertion sequence of *Pseudomonas putida* KT2440 generates different circular intermediates allowing for modular transposition.

Elena Parés-Guillén[1,2], Luis Yuste[1], Fernando Rojo[1,*], Renata Moreno[1,*]

[1] Department of Microbial Biotechnology, Centro Nacional de Biotecnología, CSIC, Madrid - 28049, Spain

[2] PhD Program in Microbiology, Sciences Faculty, Universidad Autónoma de Madrid

* To whom correspondence should be addressed. Tel: +34 91 585 4539; Fax: +34 91 585 4506;. Correspondence may also be addressed to R. Moreno. Tel: +34 91 585 4571; Fax: +34 91 585 4506;

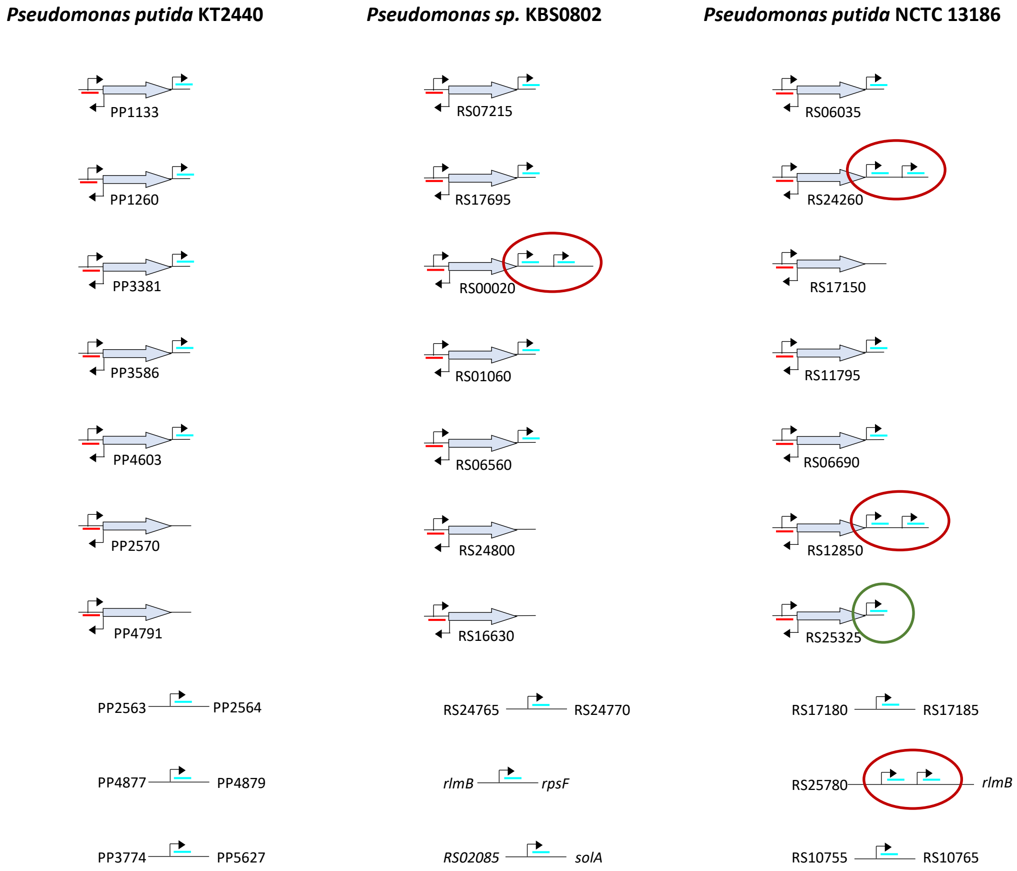

**Figure S1**. ISPpu9 and *ssr9* copies in strains *P. putida* KT2440, *Pseudomonas sp*. KBS0802 and *P. putida* NCTC13186, ordered by gene context (those in the same row are flanked by the same genes). The transposase gene (*tnp*) is indicated with grey arrows, *asr9* is in red and *ssr9* in blue. The duplicated copies of *ssr9* are highlighted with a red circle. The green circle shows a copy of *ssr9* in *P. putida* NCTC13186 that is absent in the other two strains. Thin arrowheads indicate the promoters identified in KT2440; their location in strains KBS0802 and NCTC13186 is only tentative. Adapted from Gómez-García et al. (2021), Nucleic Acids Research 49: 9211.

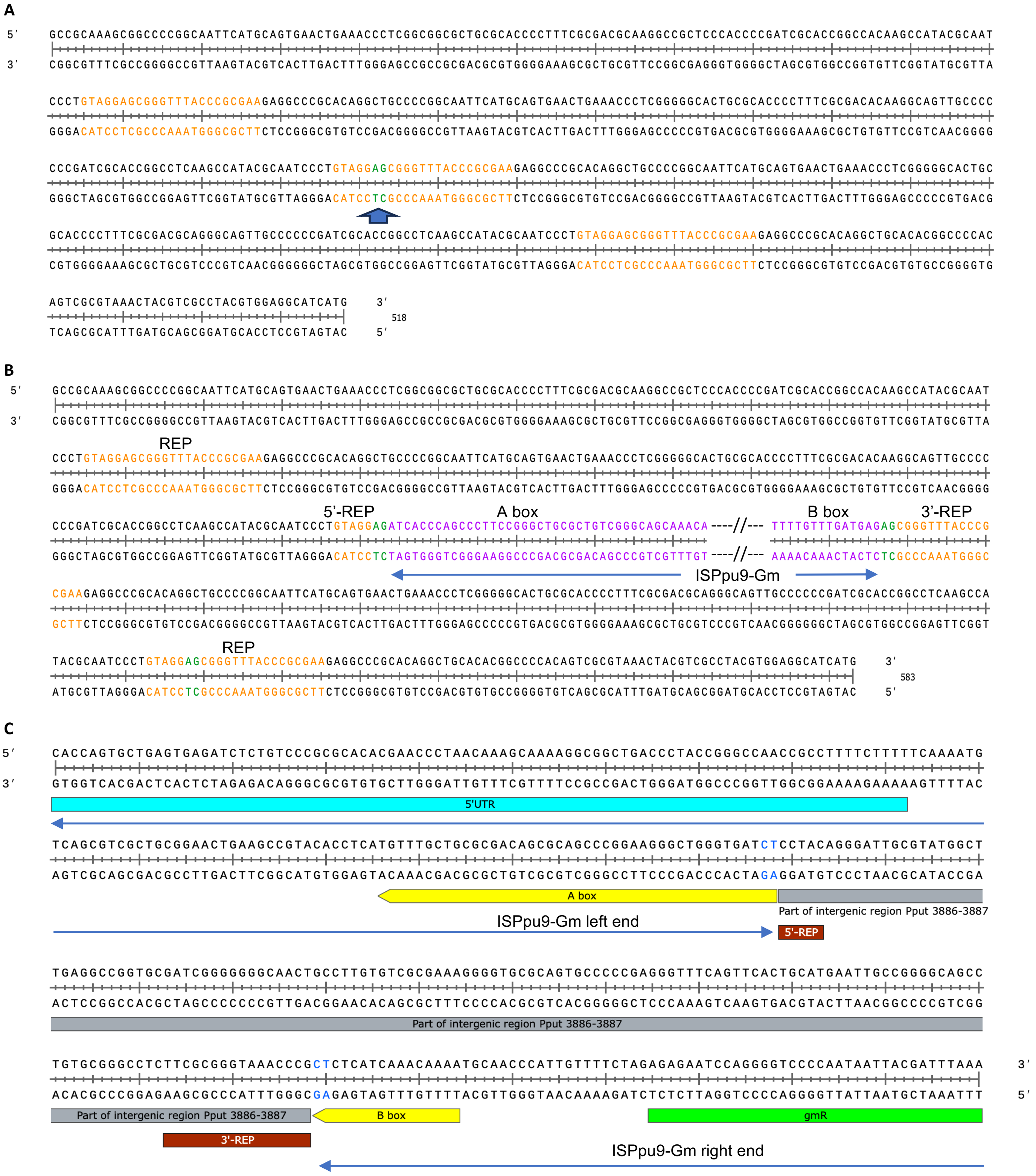

**Figure S2.** **(A)** Intergenic region at which ISPpu9-Gm is inserted in strain F1-28. The insertion site is indicated with an arrow. The three REP elements present in this region are in brown. **(B)** The left and right regions of ISPpu9-Gm inserted at the central REP element of the intergenic region shown in (A) are indicated in purple. The two AG dinucleotides that served as recombination spot in the minicircle analysed are indicated in green font. **(C)** Amplified DNA fragment corresponding to the Junc-2-type minicircle formed by ISPpu9-Gm in strain F1-28. Note that the A box, rather than recombining with the 3'-end of the B box, has recombined with the next REP element present downstream of ISPpu9-Gm, so that the intergenic region between these two REPs (in grey) lies between the 3'-end of the B box and the 5'-end of the A box. See Fig. 5B for a scheme of this minicircle.

**Table S1**. Oligonucleotides used

| Oligonucleotide sequence (5´–3´) | Name | Use |
|---|---|---|
| **Reporter fusions to *lacZ* (ISPpu9)** | | |
| gta*ggatcc*cacccagcccttcc | cmFus-D | Transcriptional fusion *Ptnp-lacZ*, forward primer |
| tgatgaagcttcgcaggataaag | PtnpR | Transcriptional fusions *Ptnp-lacZ*, *Pjunc1-lacZ* and *Pjunc2-lacZ,* reverse primer |
| acgtcagatctgcactgacat | circ-3´-2 | Transcriptional fusion *Pjunc1-lacZ*, forward primer |
| ctcgaatggataccagctgc | circ-3'tnp | Transcriptional fusion *Pjunc2-lacZ*, forward primer |
| gccgtaagcttcatgtttgctgcgcgac | BAjunc-rev | Transcriptional fusion *BAjunc-lacZ* |
| cttgaaagcttagaaaacaatgggtt | BAjunc-dir | Transcriptional fusion *BAjunc-lacZ* |
| **Reporter fusions to *lacZ* (ISPpu10)** | | |
| ggtaggaattgaagggaggaggagg | Ptnp10 dir | Transcriptional fusion *Ptnp-lacZ* (-199 to -70 rel. ATG) |
| tattgaagcttgctcgcgaggcatt | Ptnp10 rev H | Transcriptional fusion *Ptnp-lacZ* (-199 to -70 rel. ATG) |
| ggtaggaattcaagggaggaggagg | Ptnp10 large | Transcriptional fusion *Ptnp-lacZ* (-191 to -12 rel. ATG) |
| gtcgaagcttgtagcgtagtgtgact | Ptnp10 large H | Transcriptional fusion *Ptnp-lacZ* (-191 to -12 rel. ATG) |
| ggcttgaattaccagttcatcttt | Pasr10 dir | Transcriptional fusion *Pasr10-lacZ* (+69 to -67 rel.ATG) |
| gagcaagcttcaatagaagcaagcctg | Pasr10 rev H | Transcriptional fusion *Pasr10-lacZ* (+69 to -67 rel.ATG) |
| **Analysis of minicircles in ISPpu9** | | |
| ccaaatggcatcggcgtta | circ-3´-1 | Analysis of minicircles |
| acgtcagatctgcactgacat | circ-3´-2 | Analysis of minicircles |
| tccctgtcggcacagctt | circ-3´-3 | Analysis of minicircles |
| ggccgcttgagtgcttagaa | circ-3´-4 | Analysis of minicircles |
| cgcctgttttttcgctagaa | circ-3´-5 | Analysis of minicircles |
| agtgctggctacgatctctgtc | circ-5´-1 | Analysis of minicircles |
| gatgaatcctcgcaggataaa | circ-5´-2 | Analysis of minicircles |
| aaccgccttttctttttcaaaa | circ-5´-3 | Analysis of minicircles |
| cagtcggcgaatgcttttc | circ-5´-4 | Analysis of minicircles |
| tgctcccgaatgagagagtgg | circ-ssr9-F | Analysis of minicircles |
| agcagcacaccagtgcggt | circ-ssr9-R | Analysis of minicircles |
| cgtttcacgctgaatatgg | 3´-6Km | Analysis of minicircles |
| gaaccgaacaggcttatgtcaa | 3´-6Gm | Analysis of minicircles |
| acgggcagtatccaatgctg | 3'-circ10 | Analysis of minicircles |
| gcgcttgggtgttgaaagga | 5'-circ10 | Analysis of minicircles |
| gaccgccacgaaatgctctc | tnp cir 3 | Analysis of minicircles |
| aactggggtatcaactggtgg | tnp cir 4 | Analysis of minicircles |
| **qPCR assays** | | |
| tccctgtcggcacagctt | circ-3´-3 | qPCR for minicircles |
| ggccgcttgagtgcttagaa | circ-3´-4 | qPCR for minicircles |
| cgcctgttttttcgctagaa | circ-3´-5 | qPCR for minicircles |
| aaccgccttttctttttcaaaa | circ-5´-3 | qPCR for minicircles |
| cagtcggcgaatgcttttc | circ-5´-4 | qPCR for minicircles |
| ttctaccctggacctccaaca | rpoN-fwd | qPCR *rpoN* |
| ggcttgttctcggcgttgt | rpoN-rev | qPCR *rpoN* |
| **Construction of ISPpu9-∆SKm and ISPpu9-∆SGm** | | |
| ggcgtttttattgtctagaatcc | Km-D | Kanamycin amplification + XbaI site |

| | | |
|---|---|---|
| aagcagggttctagagcggaaaag | Km-R | Kanamycin amplification + XbaI site |
| gaatctagaggtccccaataattac | Gm-D | Gentamycin amplification + XbaI site |
| cgatctagagttatgcagcggaaaag | Gm-R | Gentamycin amplification + XbaI site |
| **PCR amplification of *hfq*** | | |
| aatctgtcccgcacctt | Hfq Up | Detection of KT2440 *hfq* gene |
| tgatgccttgtcaccgt | Hfq Down | Detection of KT2440 *hfq* gene |
| **PCR amplification of pKNG101 genes** | | |
| tactgtgttagatgcaatca | sacB-pKNG-dir | Detection of *sacB*, forward primer |
| agcgacaaattcgatgc | sacB-pKNG-rev | Detection of *sacB*, reverse primer |
| aatgctctctcacatcgt | Str-pKNG-dir | Detection of Sm-resistance gene, forward primer |
| gcgttgctcctcttctc | Str-pKNG-rev | Detection of Sm-resistance gene, reverse primer |
| **Sequencing of the ISPpu9-Km or ISPpu9-Gm insertion sites** | | |
| agtgctggctacgatctctgtc | circ-5´-1 | Hybridises at the 5'-end of *tnp*, reverse strand |
| cggtttgccatacagacgataa | 5'Km-seq | Hybridises at the 3'-end of Km-resistance determinant, forward strand |
| gaaccgaacaggcttatgtcaa | 5'Gm-seq | Hybridises at the 3'-end of Gm-resistance determinant, forward strand |
| **Sequencing of the pGEMT and pSEVA225 inserts** | | |
| cgccagggttttcccagtcacgac | F24 | Sequencing of pGEM-T Easy derivatives |
| agcggataacaatttcacacagga | R24 | Sequencing of pGEM-T Easy derivatives |
| **Probe for Southern blot** | | |
| gctgcgtccggtcattg | ISPpu9FwSB | Amplifies the ISPpu9 *tnp* gene |
| actaatgcagtgtccaccc | ISPpu9RevSB | Amplifies the ISPpu9 *tnp* gene |
| **Sequencing of extra *ssr9* copy in KT2440Δ*hfq* PP_4791** | | |
| cgttgctggcagggcgta | 4790dir | Detection of extra *ssr9* copy in KT2440Δ*hfq* |
| ggtggggtatcgggtgttc | 4792rev | Detection of extra *ssr9* copy in KT2440Δ*hfq* |
| **Detection of orphan ssr9 copies in KT2440Δ*hfq*** | | |
| ggtactccaggcacagcgt | 2563dir | Detection of orphan *ssr9* copy in KT2440Δ*hfq* |
| gctcgtggatgtggcctga | 2563rev | Detection of orphan *ssr9* copy in KT2440Δ*hfq* |
| ctcgctcatccatcgctg | 3774dir | Detection of orphan *ssr9* copy in KT2440Δ*hfq* |
| agcgactcggtttctagcac | 5627rev | Detection of orphan *ssr9* copy in KT2440Δ*hfq* |
| atgatttcgtaatgacgcat | 4877dir | Detection of orphan *ssr9* copy in KT2440Δ*hfq* |
| cggcaggtcaagcgctga | 4879rev | Detection of orphan *ssr9* copy in KT2440Δ*hfq* |
| **Detection of *tnp*-associated *ssr9* copies in KT2440Δ*hfq*** | | |
| aagccaagaccggcaggt | tnp-dir | Detection of *ssr9* in KT2440Δ*hfq* |
| agtggcgcgggatatgct | 1134-rev | Detection of *ssr9* in KT2440Δ*hfq* |
| ggtcgagggactggtaga | 4602-rev | Detection of *ssr9* in KT2440Δ*hfq* |
| ccgcgggacctggtgaac | ghr7-rev | Detection of *ssr9* in KT2440Δ*hfq* |
| gagccagcgacgaaggga | mdt-rev | Detection of *ssr9* in KT2440Δ*hfq* |
| cacttcactgatggtgac | ptxs-rev | Detection of *ssr9* in KT2440Δ*hfq* |

**Table S2**. A and B sequences flanking the ISPpu9 *tnp* gene, and A' and B' sequences flanking the *ssr9* gene, in *P. putida* KT2440. The AG dinucleotide present at the 5' end of the A or A' boxes, and at the 3' end of the B/B' boxes, is indicated in bold face. Positions in the A' and B' boxes that differ from those of the A or B boxes are in red. The *tnp* gene associated to each box is indicated. In the case of the *ssr9* genes, which are still not annotated at the *Pseudomonas* Genome Database, the flanking genes are specified. Mismatches relative to the A and B boxes present in PP_1133 are indicated in red.

| Sequence | Gene or intergenic region | Type |
|---|---|---|
| **AG**ATCACCCAGCCCTTCCGGGCTGCGCTGTCGCGCAGCAAACA | PP_1133 (*tnp*) | A |
| **AG**ATCACCCAGCCCTTCCGGGCTGCGCTGTCGCGCAGCAAACA | PP_1260 (*tnp*) | A |
| **AG**ATCACCCAGCCCTTCCGGGCTGCGCTGTCGCGCAGCAAACA | PP_3381 (*tnp*) | A |
| **AG**ATCACCCAGCCCTTCCGGGCTGCGCTGTCGCGCAGCAAACA | PP_3586 (*tnp*) | A |
| **AG**ATCACCCAGCCCTTCCGGGCTGCGCTGTCGCGCAGCAAACA | PP_4603 (*tnp*) | A |
| **AG**ATCACCCAGCCCTTCCGGGCTGCGCTGTCGCGCAGCAAACA | PP_2570 (*tnp*) | A |
| **AG**ATCACCCAGCCCTTCCGGGCTGCGCTGTCGCGCAGCAAACA | PP_4791 (*tnp*) | A |
| **AG**ATCACCCAGCCCTTCCGGGCTGCGCTGTCGCACAGCAAACA | PP_1133 (*ssr9*) | A' |
| **AG**ATCACCCAGCCCTTCCGGGCTGCGCTGTCGCACAGCAAACA | PP_1260 (*ssr9*) | A' |
| **AG**ATCACCCAGCCCTTCCGGGCTGCGCTGTCGCACAGCAAACA | PP_3381 (*ssr9*) | A' |
| **AG**ATCACCCAGCCCTTCCGGGCTGCGCTGTCGCACAGCAAACA | PP_3586 (*ssr9*) | A' |
| **AG**ATCACCCAGCCCTTCCGGGCTGCGCTGTCGCACAGCAAACA | PP_4603 (*ssr9*) | A' |
| **AG**ATCACCCAGCCCTTCCGGGCTGCGCTGTCGCACAGCAAACA | PP_2563-PP_2564 (*ssr9*) | A' |
| **AG**ATCACCCAGCCCTTCCGGGCTGCGCTGTCGCACAGCAAACA | *rlmB-rpsF* (*ssr9*) | A' |
| **AG**ATTACCCAGCCCTTCCAGGCTGCGCTGTCGCACAGCAAACA | PP_3774-PP_5627 (*ssr9*) | A' |
| TTTTGTTTGATGAG**AG** | PP_1133 (*tnp*) | B |
| TTTTGTTTGATGAG**AG** | PP_1260 (*tnp*) | B |
| TTTTGTTTGATGAG**AG** | PP_3381 (*tnp*) | B |
| TTTTGTTTGATGAG**AG** | PP_3586 (*tnp*) | B |
| TTTTGTTTGATGAG**AG** | PP_4603 (*tnp*) | B |
| TTTTGTTTGATGAG**AG** | PP_2570 (*tnp*) | B |
| TTTTGTTTGATGAG**AG** | PP_4791 (*tnp*) | B |
| TTTTGTTTGATGAG**AG** | *rlmB-rpsF* (*ssr9-'tnp*) | B |
| TTTTGTTTGATAGA**AG** | PP_1133 (*ssr9*) | B' |
| TTTTGTTTGATAGA**AG** | PP_1260 (*ssr9*) | B' |
| TTTTGTTTGATAGA**AG** | PP_3381 (*ssr9*) | B' |
| TTTTGTTTGATAGA**AG** | PP_3586 (*ssr9*) | B' |
| TTTTGTTTGATAGA**AG** | PP_4603 (*ssr9*) | B' |
| TTTTGTTTGATAGA**AG** | PP_2563-PP_2564 (*ssr9*) | B' |
| TTTTGTTTGATAGA**AG** | *rlmB-rpsF* (*ssr9-'tnp*) | B' |
| TTTTGTTTGATAGA**AG** | PP_3774-PP_5627 (*ssr9*) | B' |

**Table S3**. IS*110*-family transposases present in the *Pseudomonas* genomes available at the *Pseudomonas* Genome Database (version 21.1; https://www.pseudomonas.com), and showing >87% amino acid identity (over the complete protein sequence) to that encoded by *P. putida* KT2440 PP_1133 gene. The presence or absence of a sequence homologous to the Asr9 and Ssr9 sRNAs, and of A/B boxes associated to the *tnp* gene, is indicated.

| **Gene** | **Name** | **Strain** | **Tnp % Aa Ident.** | **Asr9 (% ident.)** | **Ssr9** | **Genome size** (Mb) |
|---|---|---|---|---|---|---|
| PP_1133 | IS*110* transposase | *P. putida* KT2440 | 100 | Yes (100) | yes | 6.18 |
| PP_1260 | IS*110* transposase | *P. putida* KT2440 | 100 | yes (100) | yes | 6.18 |
| PP_4603 | IS*110* transposase | *P. putida* KT2440 | 100 | yes (100) | yes | 6.18 |
| PP_4791 | IS*110* transposase | *P. putida* KT2440 | 100 | yes (100) | no | 6.18 |
| EL224_RS06035 | IS*110* transposase | *P. putida* NCTC13186 | 100 | yes (100) | yes | 6.13 |
| EL224_RS06690 | IS*110* transposase | *P. putida* NCTC13186 | 100 | yes (100) | yes | 6.13 |
| EL224_RS24260 | IS*110* transposase | *P. putida* NCTC13186 | 100 | yes (100) | yes | 6.13 |
| EL224_RS25325 | IS*110* transposase | *P. putida* NCTC13186 | 100 | yes (100) | yes | 6.13 |
| FFH79_RS25815 | IS*110* transposase | *P. sp.* KBS0802 | 100 | yes (100) | no | 6.20 |
| PP_2570 | IS*110* transposase | *P. putida* KT2440 | 99.8 | yes (100) | no | 6.18 |
| PP_3381 | IS*110* transposase | *P. putida* KT2440 | 99.8 | yes (100) | yes | 6.18 |
| PP_3586 | IS*110* transposase | *P. putida* KT2440 | 99.8 | yes (100) | yes | 6.18 |
| EL224_RS11795 | IS*110* transposase | *P. putida* NCTC13186 | 99.8 | yes (100) | yes | 6.13 |
| EL224_RS12850 | IS*110* transposase | *P. putida* NCTC13186 | 99.8 | yes (100) | yes | 6.13 |
| EL224_RS17150 | IS*110* transposase | *P. putida* NCTC13186 | 99.8 | yes (100) | no | 6.13 |
| FFH79_RS06035 | IS*110* transposase | *P. sp.* KBS0802 | 99.8 | yes (100) | yes | 6.20 |
| FFH79_RS06690 | IS*110* transposase | *P. sp.* KBS0802 | 99.8 | yes (100) | yes | 6.20 |
| FFH79_RS12195 | IS*110* transposase | *P. sp.* KBS0802 | 99.8 | yes (100) | yes | 6.20 |
| FFH79_RS13235 | IS*110* transposase | *P. sp.* KBS0802 | 99.8 | yes (100) | yes | 6.20 |
| FFH79_RS17640 | IS*110* transposase | *P. sp.* KBS0802 | 99.8 | yes (100) | no | 6.20 |
| FFH79_RS24750 | IS*110* transposase | *P. sp.* KBS0802 | 99.8 | yes (100) | yes | 6.20 |
| CHN49_RS16720 | IS*110* transposase | *P. putida* B4 | 99.1 | yes (99.4) | no | 6.01 |
| CHN49_RS16725 | IS*110* transposase | *P. putida* B4 | 99.1 | yes (99.4) | no | 6.01 |
| CHN49_RS23230 | IS*110* transposase | *P. putida* B4 | 99.1 | yes (99.4) | no | 6.01 |
| PPUBIRD1_1486 | IS*110* transposase | *P. putida* BIRD-1 | 97.5 | yes (95.6) | no | 5.73 |
| Q5O_RS07455 | IS*110* transposase | *P. putida* JB | 97.5 | yes (95.6) | no | 5.84 |
| RPPX_RS01955 | IS*110* transposase | *P. putida* S12 | 97.5 | yes (96.7) | no | 5.79 |
| RPPX_RS14455 | IS*110* transposase | *P. putida* S12 | 91.6 | yes (96.7) | no | 5.79 |
| RPPX_RS14735 | IS*110* transposase | *P. putida* S12 | 91.6 | yes (96.7) | no | 5.79 |
| RPPX_RS20870 | IS*110* transposase | *P. putida* S12 | 91.6 | yes (95.6) | no | 5.79 |
| DVB73_RS00600 | IS*110* transposase | *P. plecoglossicida* XSDHY-P | 88.9 | yes (96.1) | no | 5.52 |
| DVB73_RS01935 | IS*110* transposase | *P. plecoglossicida* XSDHY-P | 88.9 | yes (96.1) | no | 5.52 |
| DVB73_RS02495 | IS*110* transposase | *P. plecoglossicida* XSDHY-P | 88.9 | yes (96.1) | no | 5.52 |
| DVB73_RS07895 | IS*110* transposase | *P. plecoglossicida* XSDHY-P | 88.9 | yes (96.1) | no | 5.52 |
| DVB73_RS08640 | IS*110* transposase | *P. plecoglossicida* XSDHY-P | 88.9 | yes (96.1) | no | 5.52 |
| DVB73_RS09500 | IS*110* transposase | *P. plecoglossicida* XSDHY-P | 88.9 | yes (96.1) | no | 5.52 |
| DVB73_RS09740 | IS*110* transposase | *P. plecoglossicida* XSDHY-P | 88.9 | yes (96.1) | no | 5.52 |
| DVB73_RS10230 | IS*110* transposase | *P. plecoglossicida* XSDHY-P | 88.9 | yes (96.1) | no | 5.52 |
| DVB73_RS11870 | IS*110* transposase | *P. plecoglossicida* XSDHY-P | 88.9 | yes (96.1) | no | 5.52 |
| DVB73_RS11975 | IS*110* transposase | *P. plecoglossicida* XSDHY-P | 88.9 | yes (96.1) | no | 5.52 |
| DVB73_RS18425 | IS*110* transposase | *P. plecoglossicida* XSDHY-P | 88.9 | yes (96.1) | no | 5.52 |
| DVB73_RS19670 | IS*110* transposase | *P. plecoglossicida* XSDHY-P | 88.9 | yes (96.1) | no | 5.52 |
| DVB73_RS19745 | IS*110* transposase | *P. plecoglossicida* XSDHY-P | 88.9 | yes (96.1) | no | 5.52 |
| DVB73_RS19885 | IS*110* transposase | *P. plecoglossicida* XSDHY-P | 88.9 | yes (96.1) | no | 5.52 |
| DVB73_RS19945 | IS*110* transposase | *P. plecoglossicida* XSDHY-P | 88.9 | yes (96.1) | no | 5.52 |
| DVB73_RS21270 | IS*110* transposase | *P. plecoglossicida* XSDHY-P | 88.9 | yes (96.1) | no | 5.52 |
| DVB73_RS22580 | IS*110* transposase | *P. plecoglossicida* XSDHY-P | 88.9 | yes (95.5) | no | 5.52 |
| DVB73_RS23175 | IS*110* transposase | *P. plecoglossicida* XSDHY-P | 88.9 | yes (95.5) | no | 5.52 |
| DVB73_RS23210 | IS*110* transposase | *P. plecoglossicida* XSDHY-P | 88.9 | yes (95.5) | no | 5.52 |
| DW66_RS09820 | IS*110* transposase | *P. putida* DLL-E4 | 87.8 | yes (96.7) | no | 6.48 |
| PPS_4449 | IS*110* transposase | *P. putida* S16 | 87.8 | yes (96.7) | no | 5.98 |

**Table S4**. Alignment of sequences showing similarity to the A or A' boxes of ISPpu9 from *P. putida* KT2440, and that are associated to either an IS*110*-like transposase, or to a *ssr9* gene, respectively. When a strain contained >1 identical copy of the A-like box, only one of them is included in the table. A consensus is presented; the underlined nucleotides indicate an inverted repeat. Conserved nucleotides are in red font; non-conserved are in black. The adenine nucleotide that characterizes A' boxes is indicated in blue.

| **Strain** | **A-box / A'-box** | **Type** |
|---|---|---|
| *P. putida* KT2440 | AGATCACCCAGCCCTTCCGGGCTGCGCTGTCGCGCAGCAAACA | A-box |
| *P. putida* NCTC13186 | AGATCACCCAGCCCTTCCGGGCTGCGCTGTCGCGCAGCAAACA | A-box |
| *P. sp.* KBS0802 | AGATCACCCAGCCCTTCCGGGCTGCGCTGTCGCGCAGCAAACA | A-box |
| *P. putida* S12 | AGATCACCCAGCCCTTCCGGGCTGCGCTGTCGCGCAGCAAACA | A-box |
| *P. putida* S16 | AGATCACCCAGCCCTTCCGGGCTGCGCTGTCGCGCAGCAAACA | A-box |
| *P. plecoglossicida* XSDHY-P | AGATCACCCAGCCCTTTCGGGCTGCGCTGTCGCGCAGCAAACA | A-box |
| *P. putida* B4 | AGATCACCCAGCCCTTCCGGGCTGCGCTGTCGCGCAGCAAACA | A-box |
| *P. putida* BIRD-1 | AGATCACCCAGCCCTTCCGGGCTGCGCTGTCGCGCAGCAAACA | A-box |
| *P. putida* DLL-E4 | AGATCACCCAGCCCTTCCGGGCTGCGCTGTCGCGCAGCAAACA | A-box |
| *P. putida* DLL-E4 | AGATCACACAGCCCTTT-GGGCTGCGCTGTCGCGCAGAGAACA | A-box |
| *P. putida* JB | AGATCACCCAGCCCTTCCGGGCTGCGCTGTCGCGCAGCAAACA | A-box |
| *P. putida* KT2440 | AGATCACCCAGCCCTTCCGGGCTGCGCTGTCGCACAGCAAACA | A'-box |
| *P. putida* KT2440 | AGATTACCCAGCCCTTCCAGGCTGCGCTGTCGCACAGCAAACA | A'-box |
| *P. putida* NCTC13186 | AGATCACCCAGCCCTTCCGGGCTGCGCTGTCGCACAGCAAACA | A'-box |
| *P. putida* NCTC13186 | AGATTACCCAGCCCTTCCAGGCTGCGCTGTCGCACAGCAAACA | A'-box |
| *P. sp.* KBS0802 | AGATCACCCAGCCCTTCCGGGCTGCGCTGTCGCACAGCAAACA | A'-box |
| *P. sp.* KBS0802 | AGATTACCCAGCCCTTCCAGGCTGCGCTGTCGCACAGCAAACA | A'-box |
| **Consensus** | AGATCACCCAGCCCTTCCGGGCTGCGCTGTCGCGCAG<br>T T A | |

**Table S5**. Sequences showing similarity to the B or B' boxes of ISPpu9 from *P. putida* KT2440, and that are associated either to an IS*110*-like transposase (B-box), or to a *ssr9* gene (B'-box). When a strain contained >1 identical copy of the B/B'-like box, only one of them is indicated. Search was performed at the *Pseudomonas* Genome Database (version 21.1; https://www.pseudomonas.com).

| Strain | Sequence | Type | Associated A/A'- Box |
|---|---|---|---|
| *P. putida* KT2440 | TTTTGTTTGATGAG**AG** | B-box | Yes |
| *P. putida* NCTC13186 | TTTTGTTTGATGAG**AG** | B-box | Yes |
| *P. sp.* KBS0802 | TTTTGTTTGATGAG**AG** | B-box | Yes |
| *P. putida* S12 | TTTTGTTTGATGAG**AG** | B-box | Yes |
| *P. putida* S16 | TTTTGTTTGATGAG**AG** | B-box | Yes |
| *P. plecoglossicida* XSDHY-P | TTTTGTTTGATGAG**AG** | B-box | Yes |
| *P. putida* B4 | TTTTGTTTGATGAG**AG** | B-box | Yes |
| *P. putida* DLL-E4 | TTTTGTTTGATGAG**AG** | B-box | Yes |
| *P. putida* BIRD1 | TTATGTTTGAT-GA**AG** | B-box | Yes |
| *P. putida* JB | TTATGTTTGAT-GA**AG** | B-box | Yes |
| *P. putida* KT2440 | TTTTGTTTGATAGA**AG** | B'-box | Yes |
| *P. putida* KBS0802 | TTTTGTTTGATAGA**AG** | B'-box | Yes |
| *P. putida* NCTC13186 | TTTTGTTTGATAGA**AG** | B'-box | Yes |
